# Loss of perinuclear theca protein ACTRT2 causes subfertility and acrosome destabilization in mice

**DOI:** 10.64898/2026.06.05.730397

**Authors:** Andjela Kovacevic, Eva Ordziniak, Leo D. Hinterlang, Lena Arevalo, Gina E. Merges, Simon Schneider, Hubert Schorle

## Abstract

Actin-related protein T2 (ACTRT2) localizes to the perinuclear theca (PT) of male germ cells, yet its functional significance remains unclear. ACTRT2 is evolutionarily conserved and exhibits significant sequence similarity to other testis-specific actin-related proteins, with the highest conservation observed within the canonical actin core domain. We generated *Actrt2*-deficient mice which displayed male subfertility with pronounced acrosomal malformations originating during the Cap phase of acrosome biogenesis. *Actrt2*-deficient male mice showed reduced fertilization rate and poor blastocysts quality. Co-immunoprecipitation identified ACTRT2 interactions with PT proteins ACTRT1, ACTRT3, ACTL7A, ACTL9, PFN3, SPEM2 and CCIN while the interaction with CYLC1 was not detected. ACTRT2 overexpression in HEK293T cells altered cell morphology and F-actin distribution. Further, cytoskeletal regulator CFL1 was enriched in testis from *Actrt2*-deficient mice. We propose that ACTRT2 is a structural component of the PT stabilizing the acroplaxome during spermiogenesis and acrosome biogenesis by modulating actin dynamics. Finally, the high degree of sequence conservation and similarity with ACTRT1 and ACTRT3 together with their similar phenotypes when deleted, indicate that ACTRT2 shares a partial functional redundancy and compensatory capacity with other Arp proteins in testis. Taken together, these findings establish ACTRT2 as a structural regulator of sperm head architecture and male fertility in mice.

## Introduction

The proper formation of sperm in the seminiferous tubules of the testis is key to successful fertilization of an egg. Spermiogenesis comprises the morphological changes of haploid round spermatids finally resulting in polarized sperm cells with acrosome, hypercondensed DNA, species-specific head morphology and flagellum. This cellular remodeling depends on unique cytoskeletal structures like the perinuclear theca (PT), which encapsulates the developing sperm nucleus except for its caudal part at the anchoring point of the flagellum. The PT can be subdivided into an apical subacrosomal part and a caudal postacrosomal part. The subacrosomal region, also known as acroplaxome, develops first during acrosome formation and was shown to interconnect the inner acrosomal membrane with the nuclear envelope (Oko und Maravei 1994). The postacrosomal region, also known as calyx, develops at later stages during sperm head elongation and resides between the cell membrane and the nuclear envelope. The PT appears as a dense network of proteins, resistant to non-denaturating detergents and high-salt buffer extractions (Longo et al., 1987; Longo & Cook, 1991). Proteomics of PT protein extracts from mouse and bovine sperm identified 500-800 proteins with many of the corresponding genes showing testis-enriched or testis-specific expression patterns (Zhang et al. 2022a; Zhang et al. 2022b). Some of those proteins had been well molecularly characterized, including CCIN (Longo et al., 1987) multiple-band proteins CYLC1 (Hess et al., 1993) and CYLC2 (Hess et al., 1995) as well as the Actin-related proteins ACTRT1 and ACTRT2 (Heid et al., 2002). However, their functional role and their importance for male fertility are still not completely understood. The increasing number of whole-exome sequencing data from infertile men combined with the ease of generating knockout mouse models using CRISPR/Cas9 gene editing, has largely contributed to unraveling the functional impact of the PT. Recent studies have delineated that sequence variants of PT genes *ACTL7a* ((Wang et al., 2021; Xin et al., 2020; Zhou et al., 2022), *ACTL9* (Dai et al., 2021), *ACTRT1* (Zhang, Wei, Zhang, et al., 2022a), *CCIN* (Zhang, Wei, Jin, et al., 2022), *CYLC1* and *CYLC2* (Jin et al., 2024; Schneider et al., 2023) are correlated with male infertility in humans. Knockout of these genes in mice caused male sub- or infertility as well, mainly originating from defects in PT integrity and acrosome detachment from the nuclear envelope leading to partial or complete fertilization failure.

Actin-related protein T2 (ACTRT2; ARP-T2) also known as actin-related protein M2 (ARPM2) is a member of actin-related protein family specifically expressed in the testicular tissue (Heid et al., 2002). As other Actin-related proteins, ACTRT2 contains an ATPase, nucleotide binding domain which is supposed to bind and hydrolyze ATP thereby regulating the assembly and function of multiprotein complexes. Due to its actin core domain, ACTRT2 is predicted to have a structural role. Transcriptional analysis of seminal plasma extracellular vesicles (EV) revealed reduced expression of ACTRT2 in patients with oligozoospermia and non-obstructive azoospermia (NOA) (Chen et al., 2025). Furthermore, mice deficient for *Actrt2* display increased vulnerability to germ cell stress leading to death by ferroptosis (Chen et al., 2025). However, despite the previous report linking the loss of ACTRT2 with male sub-/infertility, its structural role within the complex network of PT proteins remains elusive.

We used CRISPR/Cas9-mediated gene-editing to generate an *Actrt2*-deficient mouse line. The loss of ACTRT2 resulted in detachment of the acrosome from the nuclear envelope leading to partial fertilization failure and male subfertility. Acrosomal defects originate from late round spermatids in which irregular acrosomal caps are observed. Co-immunoprecipitation experiments identified other testis-specific actin-related proteins ACTRT1, ACTRT3, ACTL7A and ACTL9 proteins as interaction partners of ACTRT2. Furthermore, ACTRT2 co-precipitated with PFN3, SPEM2 and CCIN which have been described as parts of the sperm PT. Finally, our results contribute to the expanding knowledge of the PT composition, essential for potential treatment strategies of male infertility and development of male contraceptive drugs.

## Methods

### Evolutionary analysis

The evolutionary rate of *Actrt2* was analyzed according to Lüke et al. (2016). *Actrt2* coding sequences for 39 rodent and 32 primate species, including *Loxodonta africana* and *Panthera leo* as outgroups, were obtained from NCBI GenBank and Ensembl genome browser (Tegenfeldt et al., 2025). Phylogenetic trees were built based on the NCBI taxonomy database using phyloT (https://phylot.biobyte.de). The webPRANK software was applied for codon-based phylogeny-aware alignment of orthologous gene sequences (Löytynoja & Goldman, 2010). Data were visualized using the ETE toolkit (Huerta-Cepas et al., 2016). Evolutionary rates and selective pressures were determined using codeML in PAML4.9 (Yang, 2007) based on the nonsynonymous/synonymous substitution rate ratio (ω = dN/dS) distinguishing between purifying (ω<1), neutral (ω=1) and positive selection (ω>1). Multiple null and alternative models (M) were applied, with M0 used as the baseline model. For each gene, codon frequency settings for M0 were optimized by likelihood. Likelihood ratio tests (LTRs) were used to evaluate whether alternative models describe the selective constraints within a dataset better than the null models. To determine overall evolutionary rate and selective pressures on the coding sequence across species, the M0 (one ratio) model assuming a single freely estimated ω for all branches was compared with the M0fix (fixed ratio) in which the evolutionary rate for all branches was constrained to ω=1. An LRT between M0 and M0fix was used to test deviation from neutral evolution. To assess differences in selective pressure between rodents and primates, the M0 model was compared with the MC (two ratio) model allowing independent ω estimates for each clade. To assess variation in evolutionary rate across coding sequences and identify codon sites under positive or purifying selection, LRT was performed comparing the null model M1a (nearly neutral), which excludes sites with ω>1, with the M2a (selection) model which does allows them. Codon sites were classified as purifying (Class 0: 0>ω>1), neutral (Class 1: ω = 1), or positively selected (Class 2a, M2a only: ω>1). Codons under purifying or positive selection were identified using Naïve and Bayes Empirical Bayes methods with posterior probabilities >0.95.

### Sequence alignment, conservation analysis and protein structure prediction

Multiple sequence alignment (MSA) of ACTRT1, ACTRT2, and ACTRT3 amino acid sequences was performed using Jalview with default alignment parameters (Procter et al., 2021). Sequence conservation was assessed based on residue identity and similarity and visualized using Jalview built-in coloring scheme. Conservation scores were calculated and displayed as a histogram. Consensus sequence was generated to summarize residue conservation and physicochemical similarity. Protein structures for ACTRT1, ACTRT2, and ACTRT3 were obtained from AlphaFold with the predicted local distance difference test (pLDDT) scores used as a measure of model confidence (Varadi et al., 2022). Structural alignment and visualization were performed using PyMOL (Schrödinger, LLC). The pLDDT values were mapped onto the structures using custom color thresholds: blue (pLDDT > 90), cyan (pLDDT = 70–90), yellow (pLDDT = 50–70), and orange (pLDDT < 50). Figures were rendered using PyMOL raytracing.

### Animals

Animal experiments were conducted according to German law of animal protection, with the approval of the local institutional animal care committees (Landesamt für Natur, Umwelt und Verbraucherschutz, North Rhine-Westphalia, approval ID: AZ81-02.04.2020.A259). *Actrt2*-deficient mice were generated by CRISPR/Cas9-mediated gene-editing in C57Bl/6J zygotes. Protospacer sequences are listed in Table S1. Electroporation of ribonucleoprotein (RNP) complexes into zygotes was performed using a GenePulser II electroporation device (BioRad, Feldkirchen, Germany) as published previously (Schneider et al. 2023). Recovered 2-cell embryos were cultured in G-TL medium (Vitrolife, Göteborg, Sweden) over night (37°C, 5% CO_2_) and transferred into the oviduct of pseudopregnant CB6F1 foster mice. Gene-edited alleles were separated by mating of founder animals with C57Bl/6J wildtype mice. The established mouse line was registered with Mouse Genome Informatics: B6-*Actrt2*^em1Hsc^ (MGI: 7733366). For all analyses males at the age of 8-20 weeks were used.

### Genomic DNA extraction and genotyping

Genomic DNA was isolated from ear biopsies using the HotShot extraction protocol (Truett et al., 2000). PCRs were performed using DreamTaq Green DNA Polymerase (Thermo Fisher, EP0712) according to the manufacturer’s instructions under following cycling conditions: 2 min 95°C; 35× (30 s 95°C; 30 s 60°C; 35 s 72°C); 2 min 72°C, using gene-specific primers (Table S2). PCR products were separated on 2% Agarose gel. The WT allele resulted in a 543 bp band, while the *Actrt2* deletion resulted in a 324 bp product.

### Fertility analysis and *in-vitro* fertilization

For fertility analysis, male WT, *Actrt2*^+/−^ and *Actrt2*^−/−^ mice at age of 8-18 weeks were mated with C57Bl/6J females (1:2/1:1) and the copulatory plugs were monitored daily. Plug-positive females were separated and monitored for pregnancies, followed by litter-size counts on first day post-partum. For IVF, 12-week-old C57Bl/6J WT female mice were superovulated and the oocytes were isolated. Sperm cells from three WT and three *Actrt2*^−/−^ mice were extracted from both cauda epididymides and were incubated in 200μl of FertiUp (Cosmo Bio USA, KYD-002-EX) before distributing among drops containing oocytes. The number of oocytes/embryos was recorded at 0h, 24h, 48h, 72h and 96h after fertilization. Blastocyst quality grading was performed 96h after fertilization (Hogan et al., 1986).

### Sampling

For the dissection of the reproductive organs, male mice were sacrificed at age of 8-18 weeks. The testes were fixed in Bouin or 4% PFA solution, or stored in liquid nitrogen, depending on further use. Cauda epididymides were collected in 1 mL of PBS preheated at 37°C and incisions of the cauda were performed to retrieve sperm. Immediately after extraction, sperm cells were washed in PBS, centrifuged (4000 rpm, 5 min, 4°C) and frozen or fixed with 4% PFA or Methanol and Acetic acid (3:1 v/v) solution, depending on the further use.

### Sperm morphology, motility and membrane integrity analysis

Freshly extracted epididymal sperm cells were diluted 1:40 and counted using Neubauer haemocytometer. Epididymal sperm activated in TYH medium (138 mM NaCl, 4.8 mM KCl, 2 mM CaCl 2, 1.2 mM KH 2PO4, 1 mM MgSO4, 5.6 mM glucose, 10 mM HEPES, 0.5 mM sodium pyruvate, 10 mM L-lactate, pH 7.4) was used for the motility analysis. Eosin-Nigrosin staining was performed to assess the sperm membrane integrity, as described previously (Schneider et al., 2020). For all analyses at least 100 sperm cells per animal were counted.

### Histology

Bouin fixed testis tissues were washed in 70% ethanol, paraffinized, embedded and sectioned at 3-5 µm using microtome. The sections were deparaffinized with Xylene twice for 10 min and hydrated in 100-70% descending alcohol row (5 min each). Tissue sections were incubated with Periodic acid (0.5%) for 10 min, rinsed with H_2_0 and incubated with Schiff reagent for 20 min. After staining, the sections were dehydrated in alcohol row and mounted with Entellan (Sigma-Aldrich). Slides were imaged at 40x magnification under bright field using 5500 B microscope.

### Immunofluorescence staining

For immunofluorescence staining of the acrosome with PNA-FITC Alexa Fluor 488 conjugate (Molecular Probes, Invitrogen, Waltham, MA, USA; L21409) and flagellum with MITO Tracker Red (Cell Signalling; 9082) epididymal sperm cells were fixed in 4% PFA. Staining of PT proteins was performed on methanol & acetic acid fixed sperm. For immunofluorescence staining HEK293T cells were grown on glass coverslips in a 24-well plate and fixed in 4% PFA for 15 min. All stainings were performed after permeabilization with 0.1-0.3% Triton X-100 for 10 min, blocking with Normal Horse Serum 2.5% and 5% Bovin Serum Albumin (BSA) for 30 min each. Epididymal sperm samples were incubated with 5 μg/ml PNA-FITC and 5 nM MITO Tracker Red for 1h at room temperature (RT), smeared on slides and mounted with DAPI containing medium (ROTImount FluorCare DAPI, Carl Roth; HP20.1). Bouin fixed testis tissue sections were permeabilized, blocked and washed in PBS before incubating with PNA-FITC 5 μg/ml for 1h at RT. Primary antibodies were incubated at 4°C overnight at dilutions listed in Table S3. Secondary antibodies were incubated for 1h at RT using VectaFluor Labeling Kit DyLight 488 or DyLight 594 (Vector Laboratoires, DI-1788, DI-1794). HEK293T nuclei were counterstained with 0.01 mg/ml Hoechst (bisBenzimide H 33342, Sigma-Aldrich) and coverslips were mounted on glass slides using Fluoroshield™ (F6182, Sigma-Aldrich). Sperm cells and testis sections were mounted using DAPI containing medium.

### RNA extraction and Quantitative reverse transcription-polymerase chain reaction (qRT-PCR)

Testicular RNA was extracted using TRIzol reagent according to the manufacturer`s protocol (Life Technologies, Carlsbad, USA; 15596018). RNA concentrations and purity were measured using NanoDrop ONE (Thermo Scientific). qRT-PCR was performed on ViiA 7 Real Time PCR System (Applied Biosystems) using Maxima SYBR Green qPCR Mastermix (Thermo Fisher; K0221). β-actin was used for normalization. Primer sequences are listed in table S4.

### Transmission Electron Microscopy (TEM)

Epididymal sperm cells and testicular tissue samples were fixed with 4% PFA and 2.5% glutaraldehyde in PBS overnight. The samples were washed in cacodylate buffer and incubated with 1% osmium tetroxide and 0.8% ferricyanate in cacodylate buffer for 2h. Samples were pelleted at 600 g and dehydrated in an ethanol row (30%, 50%, 70%, 90%, 95%, 100%) and propylene oxide, with addition of 0.5% uranyl acetate at the dehydration step with 70% ethanol. Finally, samples were infiltrated with propylene oxide/Epon mixtures (1:1, 1:2) followed by infiltration with pure Epon. The blocks were incubated at 60°C for 48h. Ultrathin (70 nm) sections were cut using an ultramicrotome (RMC Boeckeler Powertome) and collected on formvar-carbon-coated TEM copper slot girds. Counterstaining was performed with uranyl acetate and lead citrate. Sections were imaged with a Crossbeam 550 (Zeiss) with a retractable STEM detector, and images were acquired at 30 kV acceleration voltage with 150 pA current using SmartSEM software (Zeiss).

### Plasmid construction

To generate plasmids used in this study *Actrt2, Actrt1, Actrt3, Actl7a, Actl9, Pfn3, Spem2, Ccin* and *Cylc1* genes were amplified from C57Bl/6J mouse testis cDNA. Overhang primers introducing suitable restriction enzyme motifs are listed in Table S5. All genes were cloned in pCMV_HA-C Clontech vector (635690) and pCMV_Myc-C Clontech vector (635689). Sequencing was performed to verify the correct sequence and insertion of each plasmid.

### Protein extraction

Testicular proteins were extracted by homogenizing the whole testis without tunica albuginea in 1:10 RIPA buffer supplemented with protease inhibitors (cOmplete ULTRA Tablets, Mini, EASYpack, Roche) followed by a centrifugation for 30 min at 20 000 g, 4°C. For protein extraction, HEK293T cells were harvested in 500 μl DMEM, centrifuged for 4 min at 1400 g, 4°C and re-suspended in RIPA buffer 1:10 supplemented with protease inhibitors. Protein lysates were incubated on ice for 15 min and sonicated for 5 min using a Bioruptor (UCD-200TM-EX). For co-IP experiments proteins were extracted using M-PER™ Mammalian Protein Extraction Reagent (Thermo Scientific) supplemented with protease inhibitors according to the manufacturer’s protocol. Samples were centrifuged at 14 000 g for 10 min and supernatant was collected. Protein concentrations were measured using 96-well-plate Pierce™ BCA Protein Assay Kit (Thermo Fisher Scientific, 23225) and Bio-Rad iMARK™ plate reader.

### Co-immunoprecipitation

Co-IP experiments were performed using Pierce® HA IP kit according to the manufacturer’s protocol. Briefly, 200 μl lysate with anti-HA agarose was added to a spin column and incubated at 4°C overnight. The flow-through was collected through 10 s pulse centrifugation. The tubes were washed three times in TBS-T (0.05% Tween 20). Next, 25 μl of 2× non-reducing sample buffer were added following by incubation at 95°C for 5 min and a pulse centrifugation for 10 s. For further analysis input, flow-through and eluate were analyzed by western blotting.

### Immunoblot

Proteins were separated on 12% SDS gel with 5% stacking gel followed by transfer to PVDF membranes using Trans Blot Turbo System (Bio-Rad). Membranes were washed with 1×TBS-T (0.1% Tween20 in TBS) for 10 min and blocked with 3% BSA or 3% milk for 1h at RT. Primary antibodies were diluted in respective blocking solutions and incubated overnight at 4°C (antibody dilutions are listed in Table S3). Membranes were washed in 1×TBS-T and incubated for 1h with polyclonal goat anti-rabbit secondary antibody IgG/HRP (P044801-2; Agilent Technologies/Dako) or polyclonal rabbit anti-mouse secondary antibody IgG/HRP (P0260; Agilent Technologies/Dako) diluted 1:2000 in blocking solution. Signal was developed using WESTARNOVA2.0 chemiluminescent substrate (Cyanagen) or SuperSignal West Femto Maximum Sensitivity Substrate (34095, Thermo Scientific). Imaging was performed at ChemiDoc MP Imaging system (Bio-Rad). Uncropped images of all membranes shown are depicted in Fig. S7.

## Results

### *Actrt2* is strongly conserved across species and expressed in male germ cells

Analysis of selective pressures showed that *Actrt2* is highly conserved in rodents and primates, with a very low evolutionary rate (ω=0.079) (Fig. S1, Table S6). Although evolutionary rates differed between rodents (ω = 0.074) and primates (ω = 0.099), this difference was not biologically relevant. Additionally, 91% of codon sites were assigned to the conserved site class while 8% evolved under neutral or relaxed constraints (Fig. S1, Table S6). The distribution of evolutionary rates over the sequence indicates overall strong conservation with several peaks of relaxed constraint (Fig. S1). Multiple sequence alignment confirmed sequence conservation between ACTRT2 and other testis-specific Arps ACTRT1 and ACTRT3, revealing a highly conserved actin-like core domain (Fig. S2). Pairwise alignment showed sequence identity of 65.8% between ACTRT1 and ACTRT2, 53.2% between ACTRT1 and ACTRT3 and 43.7% between ACTRT2 and ACTRT3, with conservation predominately localized in the actin-like domain. Structural alignments of predicted protein models showed high structural similarity between ACTRT2 and ACTRT1 with low root-mean-square deviation (RMSD = 0.44 Å) and ACTRT2 and ACTRT3 (RMSD = 0.54 Å) (Fig. 1 A, B; S3 A, B). The low conservation residues were predominately surface exposed suggesting a role in functional specialization or localization rather than structural stability.

**Figure 1:**
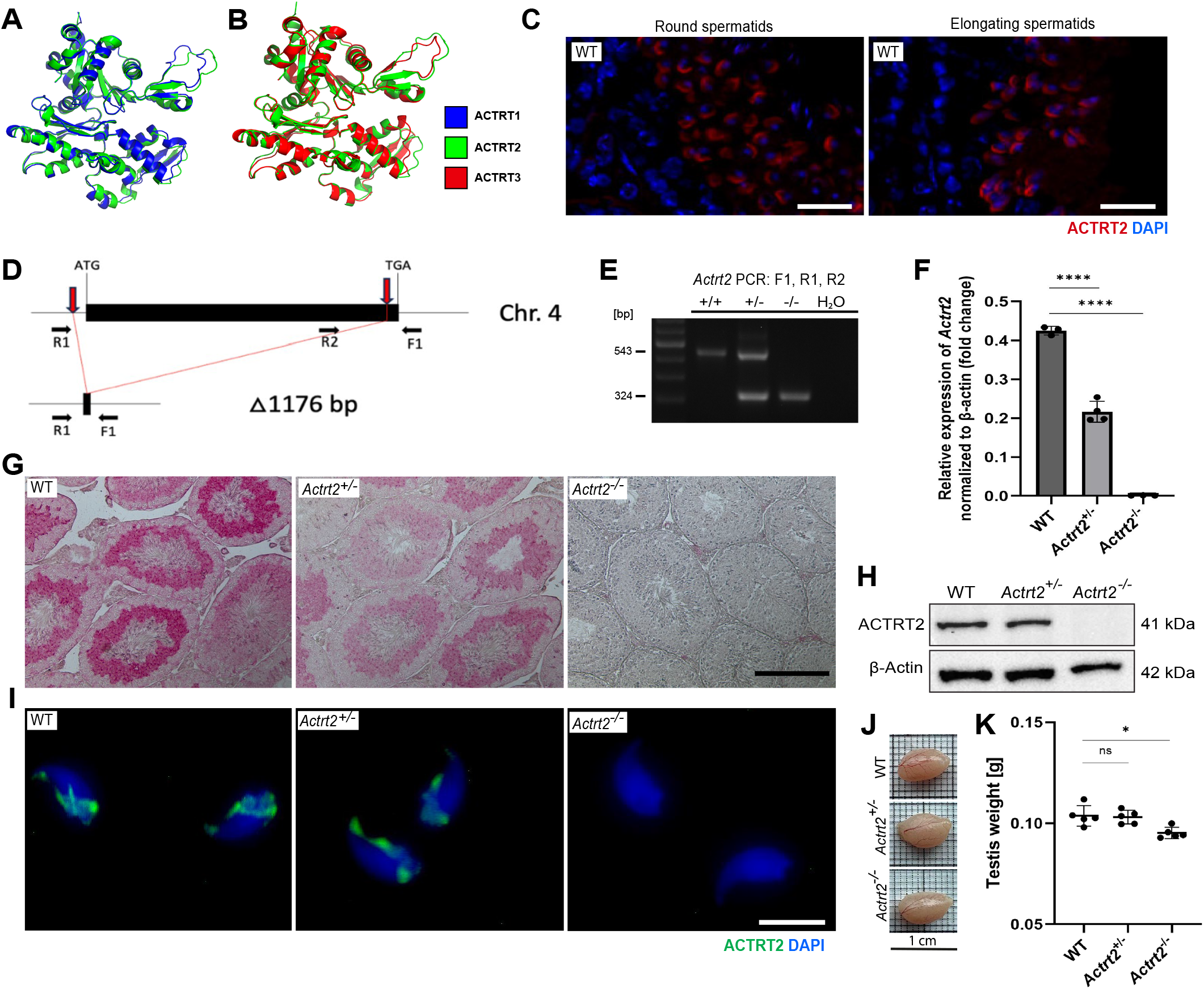
*Actrt2* is expressed in haploid male germ cells. (A) Structural alignment of ACTRT2 (green) and ACTRT1 (blue) generated using AlphaFold and visualized in PyMol. High degree of structural similarity is observed with minor deviations in loop regions. Confidence mapping is shown in Fig. S3 A. (B) Structural alignment of ACTRT2 (green) and ACTRT3 (red) generated using AlphaFold and visualized in PyMol. High degree of structural similarity is observed with minor deviations in loop regions. Confidence mapping is shown in Fig. S3 B. (C) Immunofluorescence staining against ACTRT2 (red) on testicular tissue sections from WT mice. Round spermatids and elongating spermatids are depicted showing ACTRT2 signal. Nuclei were counterstained with DAPI (blue). Scale bar: 50 μm. (D) Schematic representation of the *Actrt2* gene locus and targeting strategy for CRISPR/Cas9-mediated generation of *Actrt2-*deficient mice. Protospacer sequences are depicted by red arrows. Genotyping primer binding sites are depicted by black arrows. (E) Representative genotyping PCR of *Actrt2-*deficient mice. Band corresponding to WT *Actrt2* allele is detected at 543 bp while allele carrying *Actrt2*-deletion is detected as a band of 324 bp. (F) Relative expression of *Actrt2* in testis tissue from WT, *Actrt2*^+/−^ and *Actrt2*^−/−^ mice analyzed by qRT-PCR and normalized to β-actin. Analysis was performed using a biological replicate of 3. ****P<0.0001 (two-tailed, unpaired Student’s t-test). (G) Immunohistochemistry staining against ACTRT2 on testicular tissue sections from WT, *Actrt2*^+/−^ and *Actrt2*^−/−^ mice. Scale bar: 200 μm. (H) Western blot analysis of ACTRT2 (41 kDa) protein levels in testicular protein lysates of WT, *Actrt2*^+/−^ and *Actrt2*^−/−^ mice. β-actin was used as load control. (n=3) (I) Immunofluorescent staining against ACTRT2 (green) in epididymal sperm of WT, *Actrt2*^+/−^ and *Actrt2*^−/−^ mice. Nuclei were counterstained with DAPI (blue). Scale bar: 10 μm. (J) Comparable photographs of testis from WT, *Actrt2*^+/−^ and *Actrt2*^−/−^ mice. Scale bar: 1 cm. (K) Average testis weight (g) of adult WT, *Actrt2*^+/−^ and *Actrt2*^−/−^ mice. Black dots represent mean values obtained for each animal included in the analysis. Columns represent mean values ± s.d. *P<0.05, ns: not significant (two-tailed, unpaired Student’s t-test).

Meta analysis of available single-cell RNA-sequencing data of adult mouse testis revealed that *Actrt2* has the highest expression compared to *Actrt1* and *Actrt3* (Lukassen et al., 2018) (Fig. S3 C). Starting from late round spermatids *Actrt2* expression increased, doubling in elongated spermatids and peaking in compacted sperm (Fig. S3 C). ACTRT2 protein was detected in the perinuclear space of round and elongating spermatids from WT testis (Fig. 1 C). Similarly, data from the ‘Human testis atlas’ revealed the highest expression of *ACTRT2* compared to *ACTRT1* and *ACTRT3* (Guo et al., 2018) (Fig. S3 D, E). All three Arps are expressed starting from round spermatids, however, only *ACTRT2* displays high expression in late elongating spermatids and sperm (Fig. S3 D, E). These expression patterns of *ACTRT1-3* in mice and humans and their strong sequence similarity and conservation across species, suggest a similar role of the ARPs with overlapping functions during spermatogenesis.

### CRISPR/Cas9 gene editing resulted in deletion of *Actrt2* in mice

CRISPR/Cas9-mediated gene editing using two sgRNAs designed to delete the coding sequence of the intronless *Actrt2* gene resulted in a mouse line carrying a 1176 bp deletion (Fig. 1 D). Establishment of the line was verified by Sanger sequencing and polymerase chain reaction (PCR) was implemented for genotyping (Fig. 1 E). The expression of *Actrt2* was reduced by half in *Actrt2*^+/−^ testes compared to WT and completely diminished in *Actrt2*^−/−^ (Fig. 1 F). The loss of ACTRT2 did not alter the expression of *Actrt1* and *Actrt3* in mouse testis (Fig. S4 A). Immunohistochemistry and Western blot analysis revealed reduced ACTRT2 protein levels in *Actrt2*^+/−^ testis compared to WT, and a complete loss of ACTRT2 in *Actrt2*^−/−^ mice as expected (Fig. 1 G-H, Fig. S4 B). Finally, epididymal sperm staining showed that ACTRT2 localizes in the calyx of WT and *Actrt2*^+/−^ mice, while it was absent from *Actrt2*^−/−^ sperm (Fig. 1 I). Although a slight reduction of the testis weight was observed in *Actrt2*^−/−^ mice, the overall testis tissue architecture was not affected, with all populations of germ cells observed in all genotypes (Fig. 1 J, K; Fig. S4 C).

### *Actrt2*-deficiency causes male subfertility in mice

To investigate the role of ACTRT2, fertility assessment was performed. *Actrt2*^−/−^ mice exhibited slightly reduced pregnancy rates of 69% (17 pregnancies/25 copulatory plugs) compared to 93.3% (12/13) in *Actrt2*^+/−^ and 82.5% (14/17) in WT (Fig. 2 A). WT and *Actrt2*^+/−^ male mice obtained average litter size of 7.3 and 6.5 pups respectively, while *Actrt2*^−/−^ males were subfertile with an average litter size of 3.7 (Fig. 2 B). Slight reduction of the epididymal sperm count and membrane integrity was observed in *Actrt2*^−/−^ mice compared to WT or *Actrt2*^+/−^ (Fig. 2 C, D, Fig. S5 A). Interestingly, sperm motility upon activation in TYH medium was not significantly altered (Fig. 2 E).

**Figure 2:**
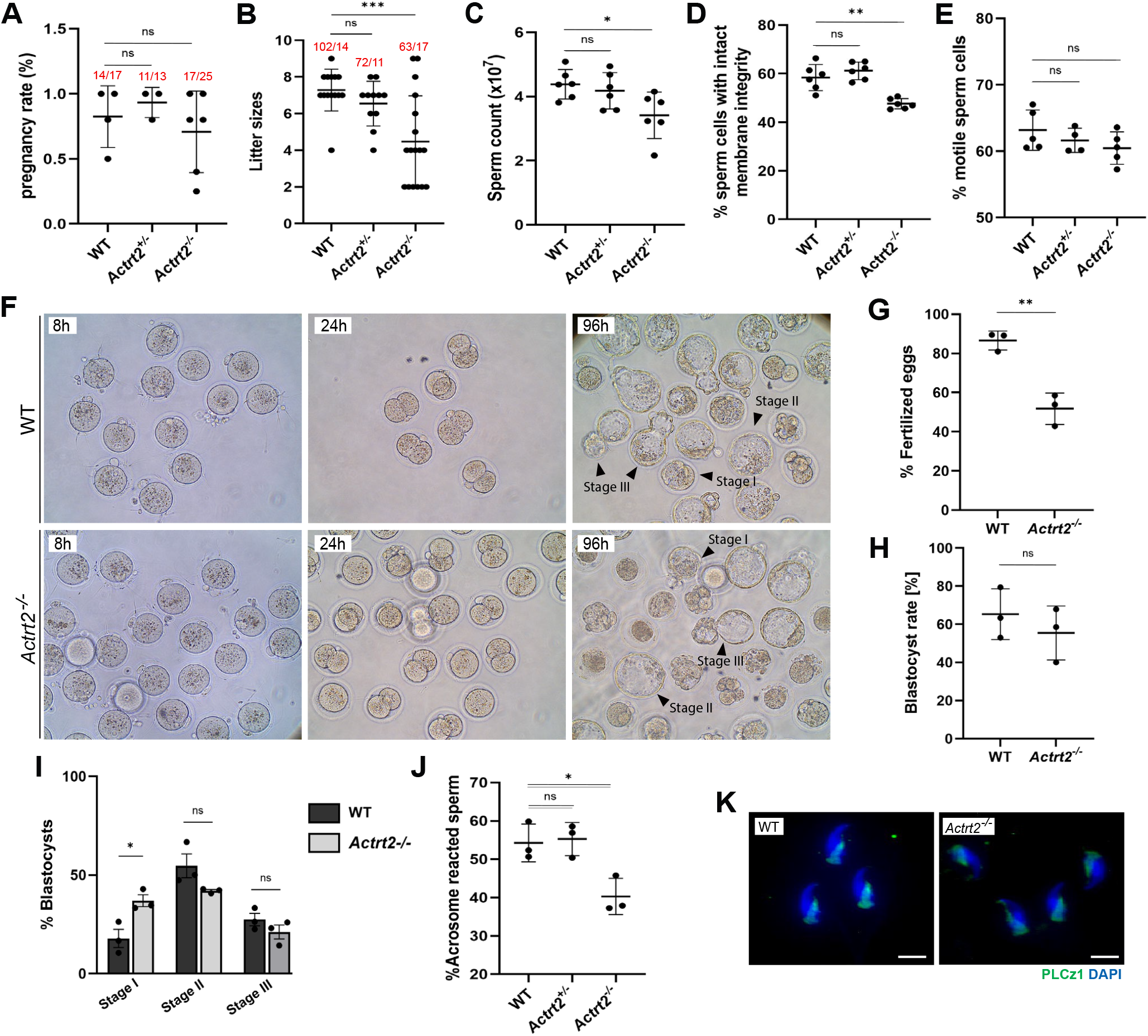
Loss of *Actrt2* disrupts male fertility in mice. (A) Pregnancy rates (%) obtained in matings of WT female with WT (n=4), *Actrt2*^+/−^ (n=3) and *Actrt2*^−/−^ (n=6) male mice. Black dots represent mean values obtained for each male included in the mating. Columns represent mean values ± s.d. Total number of pregnancies per total number of copulatory plugs observed are depicted above each bar. ns: not significant (two-tailed, unpaired Student’s t-test). (L) Total litter sizes observed during fertility assessment of WT (n=4), *Actrt2*^+/−^ (n=3) and *Actrt2*^−/−^ (n=6) male mice. Black dots represent each litter count on day 1 postpartum. Columns represent mean values ± s.d. Total number of pups born per number of litters observed are depicted above each bar. ***P<0.001, ns: not significant (Welch’s t-test). (B) Epididymal sperm count (×10^7^) of WT, *Actrt2*^+/−^ and *Actrt2*^−/−^ male mice (n=6 for each genotype). Black dots represent mean values obtained for each animal included in the analysis. Columns represent mean values ± s.d. *P<0.05, ns: not significant (two-tailed, unpaired Student’s t-test). (C) Quantification of sperm cells with intact membrane integrity from WT, *Actrt2*^+/−^ and *Actrt2*^−/−^ male mice analyzed by Eosin-Nigrosine staining (n=6 for each genotype). Black dots represent mean values obtained for each animal included in the analysis. Columns represent mean values ± s.d. A minimum of 100 cells per sample was counted. **P<0.005, ns: not significant (two-tailed, unpaired Student’s t-test). (D) Quantification of motile epididymal sperm activated in THY medium from WT (n=5), *Actrt2*^+/−^ (n=4) and *Actrt2*^−/−^ (n=5). Black dots represent mean values obtained for each animal included in the analysis. Columns represent mean values ± s.d. ns: not significant (two-tailed, unpaired Student’s t-test). (E) Representative images of the IVF assay performed using sperm from WT and *Actrt2*^−/−^ mice at 8, 24 and 96 h after fertilization. Black arrowheads depict blastocyst stages I-III. (F) Percentage of oocytes fertilized by WT and *Actrt2*^−/−^ sperm in IVF. Oocytes displaying visible maternal and paternal pronuclei were counted as successfully fertilized. Black dots represent mean values obtained for each animal included in the analysis. Columns represent mean values ± s.d. (n=3). **P<0.005 (two-tailed, unpaired Student’s t-test). (G) Percentage of fertilized oocyte which developed into blastocysts after 96h of incubation. Black dots represent mean values obtained for each animal included in the analysis. Columns represent mean values ± s.d. (n=3). ns: not significant (two-tailed, unpaired Student’s t-test). (H) Percentage of blastocysts at stage I, stage II and stage III obtained by IVF with WT and *Actrt2*^−/−^ sperm. Black dots represent mean values obtained for each animal included in the analysis. Columns represent mean values ± s.d. (n=3). *P<0.05, ns: not significant (two-tailed, unpaired Student’s t-test). (I) Quantification of acrosome reacted sperm cells from WT, *Actrt2*^+/−^ and *Actrt2*^−/−^ male mice upon capacitation and incubation with ionophore A23187 (n=3). Black dots represent mean values obtained for each animal included in the analysis. Columns represent mean values ± s.d. *P<0.05, ns: not significant (two-tailed, unpaired Student’s t-test). (J) Immunofluorescence staining against PLC ζ1 (green) on epididymal sperm from WT and *Actrt2*^−/−^ mice. Nuclei were counterstained with DAPI (blue). Scale bar: 10 μm. (n=3)

*In-vitro* fertilization of WT oocytes was performed to evaluate the role of ACTRT2 in fertilization. After 8h of incubation, oocytes displaying visible pronuclei were counted revealing a reduced fertilization rate of 51% in *Actrt2*^−/−^ mice compared to 86% in WT (Fig. 2 F, G). After 96h blastocyst formation was modestly reduced in *Actrt2*^−/−^ mice (55%) while the blastocyst rate in WT was 65% (Fig. 2 F, H). Next, the quality and developmental stage of blastocysts were evaluated (Hogan et al., 1986). While stage I blastocysts were increased in *Actrt2*^−/−^ (37%) compared to WT (18%), fully expanded, stage II blastocysts accounted for 45% in WT and in 42% *Actrt2*^−/−^ embryos. Stage III (hatching) blastocysts were slightly decreased in *Actrt2*^−/−^ embryos (21% vs 27% in WT) (Fig. 2 F, I). The acrosome reaction assessment revealed that roughly 55% of sperm cells from WT and *Actrt2*^+/−^ mice underwent acrosome reaction upon capacitation and incubation with ionophore A23187, compared to only 40% in *Actrt2*^−/−^ sperm, indicating impaired acrosome function underlying the reduced fertilization rate (Fig. 2 J, Fig. S5 B). Phospholipase C zeta 1 (PLCζ1) required for oocyte activation, remained correctly localized in the calyx of both WT and *Actrt2*^−/−^ sperm (Fig. 2 K) (Yoon & Fissore, 2007).

### *Actrt2*^−/−^ sperm cells display acrosome defects

To investigate acrosome structure in detail, morphological analysis of epididymal sperm was performed. While in WT and *Actrt2*^+/−^ sperm acrosomes displayed as cap-like structures on the apical portion of the sperm head, in *Actrt2*^−/−^ mice 50% of acrosomes appeared misshapen or shortened (Fig. 3 A, B). Protein levels of the acrosomal marker Sperm Acrosome Associated 1 (SPACA1) were reduced in testicular protein lysates from *Actrt2*^+/−^ and *Actrt2*^−/−^ mice compared to WT (Fig. 3 C, Fig. S5 D). Sperm flagella stained with MITO-red appeared unaltered in all three genotypes (Fig. 3 A, Fig. S5 C). In Transmission electron micrographs (TEM) of epididymal sperm from WT and *Actrt2*^+/−^ mice we observed electron-dense nuclei tightly surrounded by PT and intact acrosomes (Fig. 3 D). The *Actrt2*^−/−^ sperm displayed electron-dense nuclei indicating that the loss of ACTRT2 does not affect chromatin compaction, however improperly evicted excess cytoplasm was observed (Fig. 3 D, yellow arrowheads). In *Actrt2*^−/−^ sperm acrosomes were detached from the nuclear envelope and shrunk towards the apical portion of the cell (Fig. 3 D, green arrowheads). The PT appeared loosened and detached from the nuclear envelope at the equatorial ring (Fig. 3 D). The axonemal structure of the sperm flagella remained intact upon loss of ACTRT2, displaying the canonical 9+2 microtubule arrangement surrounded by a mitochondrial sheath and outer dense fibers in the midpiece (Fig. 3 E).

**Figure 3:**
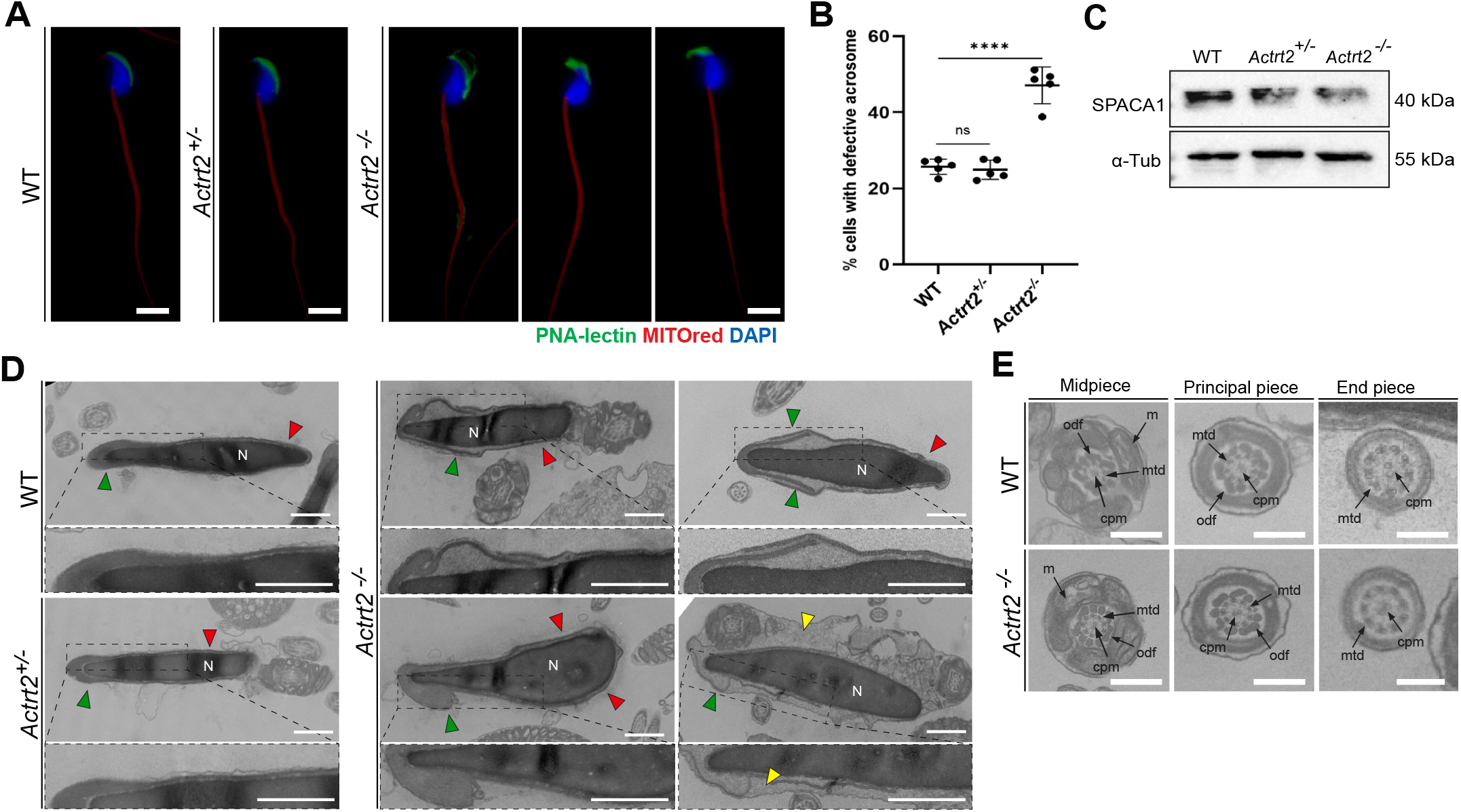
Loss of ACTRT2 causes acrosome defects in epididymal sperm. (A) Immunofluorescence staining of epididymal sperm from WT, *Actrt2*^+/−^ and *Actrt2*^−/−^ male mice with acrosomal marker PNA-lectin (green) and mitochondrial marker MITOred (red). Nuclei were counterstained with DAPI (blue). Representative images of different acrosome defects in *Actrt2*^−/−^ sperm are depicted. Scale bar: 5 μm. (B) Quantification of sperm cells with acrosome abnormalities observed by PNA-lectin staining in WT, *Actrt2*^+/−^ and *Actrt2*^−/−^ male mice. Black dots represent mean values obtained for each animal included in the analysis. Columns represent mean values ± s.d. (n=5). ****P<0.0001 (two-tailed, unpaired Student’s t-test). (C) Western blot analysis of SPACA1 protein levels in testicular protein lysates of WT, *Actrt2*^+/−^ and *Actrt2*^−/−^ mice. α-Tubulin was used as load control. (n=3) (D) TEM micrographs of epididymal sperm from WT, *Actrt2*^+/−^ and *Actrt2*^−/−^ mice. Green arrowheads depict acrosomes, and red arrowheads show calyx region of the PT. Sperm cells from *Actrt2*^−/−^ display defective acrosomes detached from the nuclear envelope. Incorrectly evicted cytoplasm is depicted with yellow arrowheads. N, nucleus. Scale bars: 1 μm. (E) TEM micrographs depicting cross sections of midpiece, principal piece and end piece of sperm flagella from WT, *Actrt2*^+/−^ and *Actrt2*^−/−^ mice. All typical structures are depicted: m, mitochondria; mtd, microtubule doublets; cpm, central-pair microtubule; odf, outer-dense fibers. Scale bar: 100 nm.

PNA-lectin staining of testicular tissue was performed to investigate the origin of acrosomal defects during spermiogenesis. Spermatids at Golgi phase of acrosome development displayed no significant differences between WT, *Actrt2*^+/−^ and *Actrt2*^−/−^ mice, having intact proacrosomal granules (Fig. 4 A). First defects of the acrosome development in *Actrt2*^−/−^ testis can be observed in late round spermatids at cap phase which displayed irregular or vacuolized acrosomal structures in contrast to smooth cap-like acrosomes found in WT and *Actrt2*^+/−^ mice (Fig. 4 A). Further defects of the acrosome such as detachment from the nucleus and accumulation at the tip of the cell were observed in the elongating spermatids in acrosome phase reflecting the malformations observed in mature sperm. TEM analysis of testicular tissue showed normal morphology of Golgi phase round spermatids in *Actrt2*-deficient mice, whereas late round spermatids exhibited a shortened acrosomal cap compared to WT (Fig. 4 B). Although the manchette formed normally in *Actrt2*^−/−^ elongating spermatids, an enlarged gap between the acrosome and manchette at the equatorial ring was observed (Fig. 4 B). A slight decrease of the Golgi-associated PDZ- and coiled-coil motif-containing protein (GOPC) required for trans-Golgi vesicle trafficking during acrosome development was observed in testicular lysates from *Actrt2*^−/−^ mice (Fig. 4 C, Fig. S5 E). Furthermore, Testicular Trans-Golgi Network 46 (TGN46) marker did not appear affected indicating that the loss of ACTRT2 does not cause severe Golgi aberrations (Fig. 4 C, Fig. S5 F). Taken together, the loss of ACTRT2 causes acrosomal defects at the late round spermatid stage while Golgi vesicle trafficking remains unaffected.

**Figure 4:**
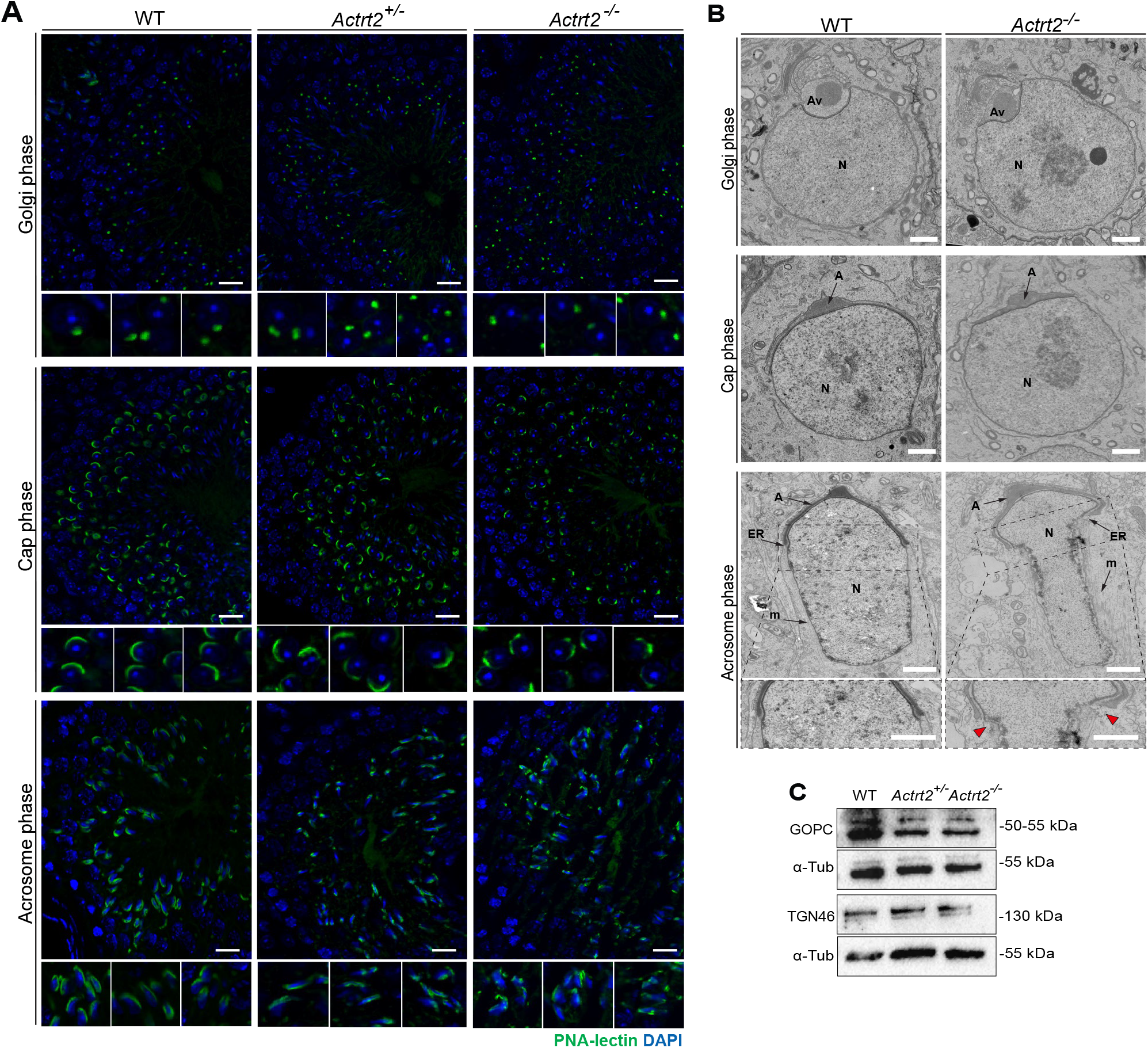
*Actrt2-*deficiency causes aberrant acrosome development during spermiogenesis. (A) Immunofluorescence staining of testicular tissue sections from WT, *Actrt2*^+/−^ and *Actrt2*^−/−^ male mice with acrosomal marker PNA-lectin (green). Nuclei were counterstained with DAPI (blue). Representative images of spermatids in Golgi phase (top panel), cap phase (middle panel) and acrosome phase (bottom panel) are shown. Inserts depict individual spermatids present in the main image at larger magnification. Staining was performed on three animals of each genotype. Scale bar: 20 μm. (B) TEM micrographs of WT and *Actrt2*^−/−^ spermatids in Golgi, cap and acrosome phase of acrosome development. In Golgi phase, acrosomal vesicles are regular in both genotypes (top panel). Spermatids in cap phase from *Actrt2*^−/−^ mice display shorter acrosomal cap compared to WT spermatids (middle panel). In acrosome phase of *Actrt2*^−/−^ mice a gap is observed between the acrosome and the manchette at the equatorial ring (bottom panel, red arrowhead). Inserts depict equatorial ring area at higher magnification. N, nucleus; Av, acrosomal vesicle; A, acrosome; m, manchette; ER, equatorial ring. Scale bar: 1 μm.

### ACTRT2 is part of PT molecular complex

Molecular interactions between ACTRT2 and other components of the PT were investigated by *in-vitro* co-immunoprecipitation (Co-IP) analysis. Expression plasmids of PT-specific proteins carrying HA- or Myc-tag were generated and transiently transfected in HEK293T cells (Fig. S5 G, H). ACTRT2 has been previously identified as interaction partner of ACTRT1 (Zhang, Wei, Zhang, et al., 2022a) and ACTRT3 (Kovacevic et al., 2026), which was here confirmed (Fig. 5 A, Fig. S6 A). Interestingly, *Actrt2*-deficiency caused drastic reduction of ACTRT1 in the calyx region of the PT of epididymal sperm and significantly diminished protein levels in testis lysates (Fig. 5 B, C, red arrowheads, Fig. S6 B). Levels of ACTRT3 appeared less reduced and its localization within the calyx was unaltered in *Actrt2*^−/−^ sperm (Fig. 5 B, D, Fig. S6 C). Interestingly, ACTRT2 co-precipitated with PFN3 and SPEM2 which are known interaction partners of ACTRT3, suggesting an overlapping role between the Arps (Fig. 5 A, Fig. S6 A) (Kovacevic et al., 2025; Li et al., 2024). Next, ACTRT2 co-precipitated with ACTL7A and ACTL9, major components of the PT required for male fertility in humans and mice. In *Actrt2*^−/−^ epididymal sperm ACTL7A appeared confined to the calyx region and diminished from the apical portion of the sperm head while it was localized in postacrosomal and subacrosomal regions of the PT of WT and *Actrt2*^+/−^ sperm (Fig. 5 B, yellow arrowhead). A strong Co-IP signal was found for CCIN - a structural PT protein required for the sperm head shaping (Fig. 5 A, Fig. S6 A). ACTRT2 co-precipitated with CCIN, but not with its interaction partner CYLC1 indicating that CCIN might be the connector between ACTRT2 and CYLC1 (Fig. 5 A, Fig. S6 A) (Zhang, Wei, Jin, et al., 2022). CCIN and CYLC1 maintained their localization in the calyx of *Actrt2*^−/−^ sperm indicating that ACTRT2 is not essential for maintenance of the calyx structure (Fig. 5 E).

**Figure 5:**
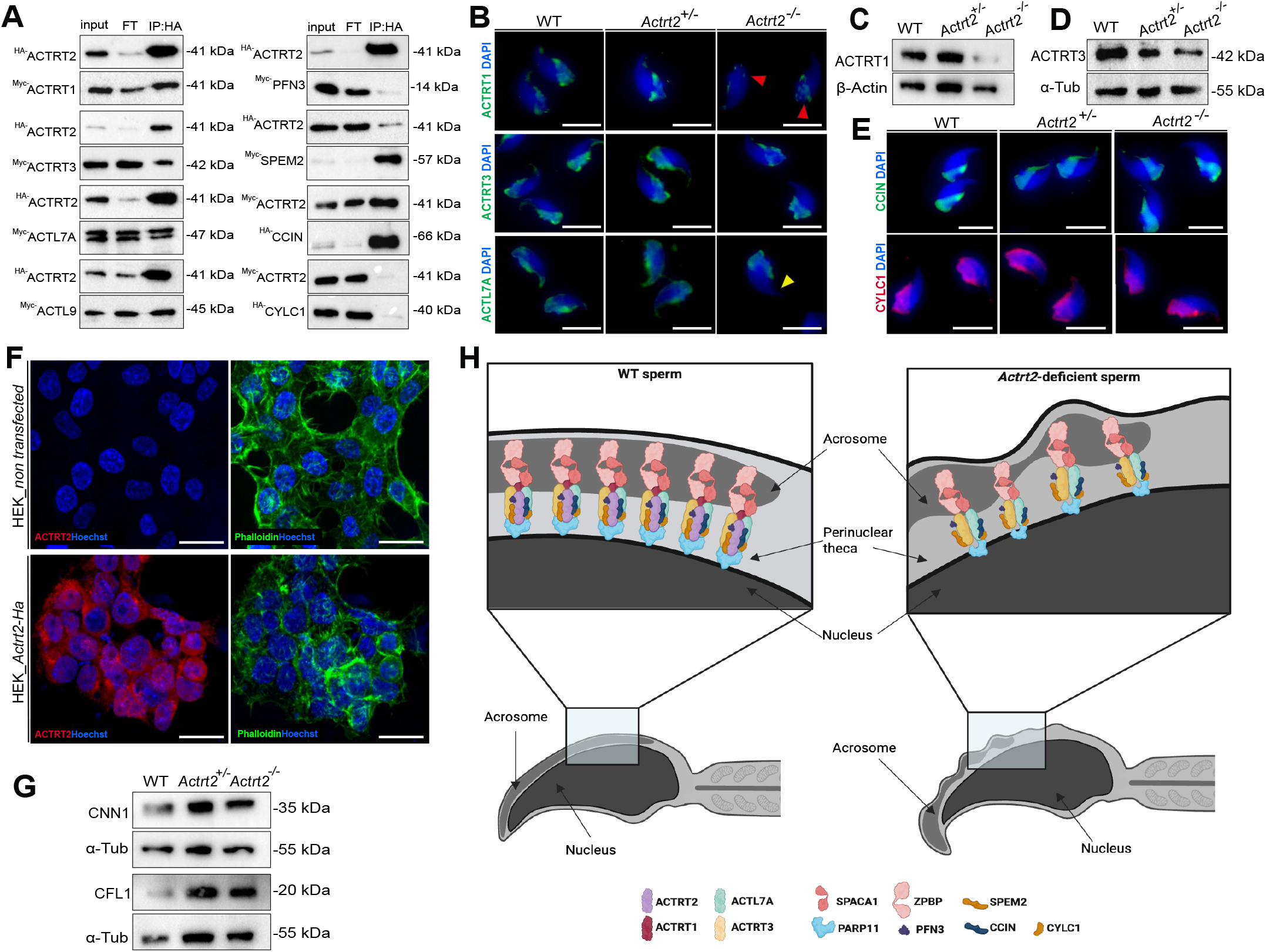
*ACTRT2* interacts with other components of the PT. (A) Co-IP analysis of proteins expressed in HEK293T cells. ACTRT2 co-precipitated with ACTRT1, ACTRT3, ACTL7A, ACTL9, PFN3, SPEM2 and CCIN. No interaction was detected between ACTRT2 and CYLC1. (B) Immunofluorescence staining against ACTRT1, ACTRT3 and ACTL7A (green) on epididymal sperm from WT, *Actrt2*^+/−^ and *Actrt2*^−/−^ mice. Red arrowheads depict reduction of ACTRT1 signal in the calyx of *Actrt2*^−/−^ sperm. Yellow arrowheads show loss of ACTL7A in the apical portion of the *Actrt2*^−/−^ sperm heads. Nuclei were counterstained with DAPI (blue). Scale bar: 10 μm. (C) Western blot analysis of ACTRT1 (42 kDa) protein levels in testicular protein lysates of WT, *Actrt2*^+/−^ and *Actrt2*^−/−^ mice. β-actin was used as load control. (n=3) (D) Western blot analysis of ACTRT3 (42 kDa) protein levels in testicular protein lysates of WT, *Actrt2*^+/−^ and *Actrt2*^−/−^ mice. α-Tubulin was used as load control. (n=3) (E) Immunofluorescence staining against CCIN (green) and CYLC1 (red) on epididymal sperm from WT, *Actrt2*^+/−^ and *Actrt2*^−/−^ mice. Nuclei were counterstained with DAPI (blue). Scale bar: 10 μm. (F) Immunofluorescence staining of F-actin marked by Phalloidin (green) and ACTRT2 (red) in non-transfected HEK239T cells (upper panel) and *Actrt2*-HA HEK239T cells (lower panel). Nuclei were counterstained with Hoechst (blue). Scale bar: 10 μm. (G) Western blot analysis of CNN1 (35 kDa) and CFL1 (20 kDa) protein levels in testicular protein lysates of WT, *Actrt2*^+/−^ and *Actrt2*^−/−^ mice. α-Tubulin was used as load control. (n=3) (H) Schematic illustration of PT complex in WT and *Actrt2*^−/−^ epididymal sperm. Destabilized PT structure is shown in *Actrt2*^−/−^ sperm, causing acrosome abnromalities and detachment from the nucleus. Created in BioRender by Schorle, H., 2026.

Phalloidin staining of non-transfected HEK239T cells revealed healthy F-actin filaments distributed throughout the cytoplasm. HEK239T cells overexpressing *Actrt2* appeared rounder, with thicker F-actin filament accumulation in the cortical region of the cell (Fig. 5 F). *Actrt3*-deficient male mice displayed increased protein levels of actin-binding cytoskeletal regulator Calponin 1 (CNN1) and actin filament turnover regulator Cofilin 1 (CFL1) suggesting the abnormal actin dynamics upon loss of ACTRT3 (Kovacevic et al., 2026). Similarly, enrichment of CNN1 and CFL1 in testicular protein lysates from *Actrt2*-deficient mice was observed, supporting the role of ACTRT2 in cytoskeletal regulation of male germ cells (Fig. 5 G, Fig. S6 D, E).

## Discussion

*Actrt2* is a highly conserved member of the Arp protein family exhibiting testis-specific expression enriched in the male germline. Here, we showed that *Actrt2*-deficient male mice are subfertile, with acrosome abnormalities and detachment from the nuclear envelope starting from round spermatids. Furthermore, ACTRT2 is an integral part of PT structural complex as shown by the interaction with ACTRT1, ACTRT3, ACTL7A, ACTL9, PFN3, SPEM2 and CCIN. HEK239T cells overexpressing *Actrt2* exhibit morphological changes demonstrating the role of ACTRT2 in actin dynamics.

*Actrt2*-knockout mouse model exhibiting severely impaired spermatogenesis accompanied by shrunken seminiferous tubules has been previously reported; however, male fertility and protein ACTRT2 levels were not assessed (Chen et al., 2025). Our results show that despite the slightly reduced testis weight in *Actrt2*-knockout mouse, seminiferous tubules were found intact, with all stages of spermatogenesis observed, leading to reduced epididymal sperm counts but not complete spermatogenesis arrest. The difference in observed results could not be explained by a different animal strain as in both studies C57BL/6 mice were used.

Recent studies demonstrated that ACTRT1, ACTRT2, ACTRT3, ACTL7A and ACTL9 interact among each other forming a sperm-specific Arp complex functioning as a core of the PT that anchors the inner acrosomal membrane (IAM) to the nuclear envelope (NE) (Fig. 5 H). ACTRT1 and ACTL7A represent central components of the IAM-NE scaffold interacting with PARP11 which is involved in nuclear envelope organization as well as with IAM protein SPACA1 (Zhang, Wei, Zhang, et al., 2022b). Here we demonstrated the interaction between ACTRT2 and CCIN which forms a complex with ACTL7A, CYLC1 and PARP11 to additionally strengthen the IAM-NE connection (Zhang, Wei, Jin, et al., 2022). Interestingly, *Ccin*-knockout mice exhibit nuclear subsidence rather than acrosomal detachment observed in Arp-deficient models indicating that it might form an additional complex with the NE (Zhang, Wei, Jin, et al., 2022).

High degree of evolutionary conservation of *Actrt2* across rodents and primates, as well as its significant sequence similarities with ACTRT1 and ACTRT3 strongly support a model in which ACTRT1-3 might exhibit partially redundant functions which might lead to cross-compensatory effects. ACTRT1 has been described as a component of the sperm PT involved in acrosomal attachment and head shaping (Zhang, Wei, Zhang, et al., 2022b), while ACTRT3 regulates actin dynamics and vesicle trafficking during acrosome biogenesis in mice (Kovacevic et al., 2026). *Actrt1*- and *Actrt3*-deficient mice exhibit male subfertility and fertilization failure due to acrosome malformations and detachment from the nuclear envelope similarly to *Actrt2*-deficient mice (Kovacevic et al., 2026; Zhang, Wei, Zhang, et al., 2022b). *Actrt1*-knockout spermatids exhibit disruption of the acroplaxome, the F-actin scaffold located between the nucleus and the acrosome. In contrast, *Actrt1*-deficiency caused manchette disorganization, which was not observed in *Actrt2*-deficient mice, indicating that ACTRT2 functions exclusively at the acroplaxome level (Zhang, Wei, Zhang, et al., 2022b). Furthermore, we showed that ACTRT2 interacts with PFN3 and SPEM2 which have been identified as interaction partners of ACTRT3 (Kovacevic et al., 2026). PFN3 is a small actin-regulating protein required for Golgi trafficking and autophagic flux in murine spermatids while SPEM2 has a role in sperm individualization and cytoplasm eviction being both essential for male fertility in mice (Li et al., 2024; Umer et al., 2021). While in *Actrt3*-knockout mice *cis*- and *trans*-Golgi trafficking were significantly affected leading to abnormal acrosome development, only slight reduction of the *trans*-Golgi marker GOPC was observed in *Actrt2*-deficient testis (Kovacevic et al., 2026). GOPC mediates transport of proacrosomal vesicles from the *trans*-Golgi network to the developing acrosome and its loss leads to acrosome malformations in early round spermatids and male infertility in mice (Yao et al., 2002). However, unaltered protein abundance of *trans*-Golgi marker TGN46 which is involved in vesicle biogenesis along with intact Golgi structure observed in TEM indicate that the *Actrt2*-deficiency did not disrupt acrosome biogenesis at Golgi phase. This distinction between the roles of ACTRT1-3 is supported by the transcriptomics data demonstrating higher expression of *Actrt2* in later stages of spermiogenesis suggesting that it’s required for maintaining the acroplaxome integrity and acrosome attachment to the nucleus rather than the initial stages of acrosome biogenesis. Alterations of the cell morphology upon overexpression of *Actrt2* and the enrichment of CNN1 and CFL1 in *Actrt2*-deficient testis, similarly to what was previously observed in Actrt3-knockout mice, demonstrate the role of ACTRT2 as a cytoskeletal regulator (Kovacevic et al., 2026). During spermiogenesis, ACTRT2 is likely involved in acrosome development and cytoplasmatic eviction via its interaction with PFN3 and SPEM2 and by regulating actin filament turnover. The loss of ACTRT2 leads to depletion of ACTRT1 from the PT resulting in the destabilization of IAM-NE scaffold and detachment of the acrosome in roughly half of the sperm cells, which could partially be rescued by ACTRT3 and ACTL7A (Fig. 5 H).

Our results demonstrate the distinct but partially redundant role of ACTRT2 in acrosome biogenesis and sperm structural integrity, similar to the functions of ACTRT1 and ACTRT3. Studies in compound *Actrt1/Actrt2/Actrt3*-knockout mouse models could unravel the degree of functional redundancy and compensatory roles of different Arps.

## Acknowledgment

We are grateful to Gaby Beine, Angela Egert, Andrea Jäger and Greta Zech for excellent technical assistance. We would like to thank the Core Facility for Microscopy of the Medical Faculty at the University of Bonn for providing support and instrumentation funded by the Deutsche Forschungsgemeinschaft (DFG, German Research Foundation, project number: 388169927).

## Author Contributions

A.K., S.S. and H.S. conceived and designed the experiments. S.S. generated *Actrt2*-deficient mouse model. A.K., E.O. and L.D.H. analyzed *Actrt2*-deficient mice. L.A. performed evolutionary analysis. G.E.M, S.S. and H.S. supervised the study. A.K., S.S. and H.S. were major contributors in writing the manuscript. All authors read and approved the final manuscript.

