## Supplements for "Loss of perinuclear theca protein ACTRT2 causes subfertility and acrosome destabilization in mice"

#### Figure S1

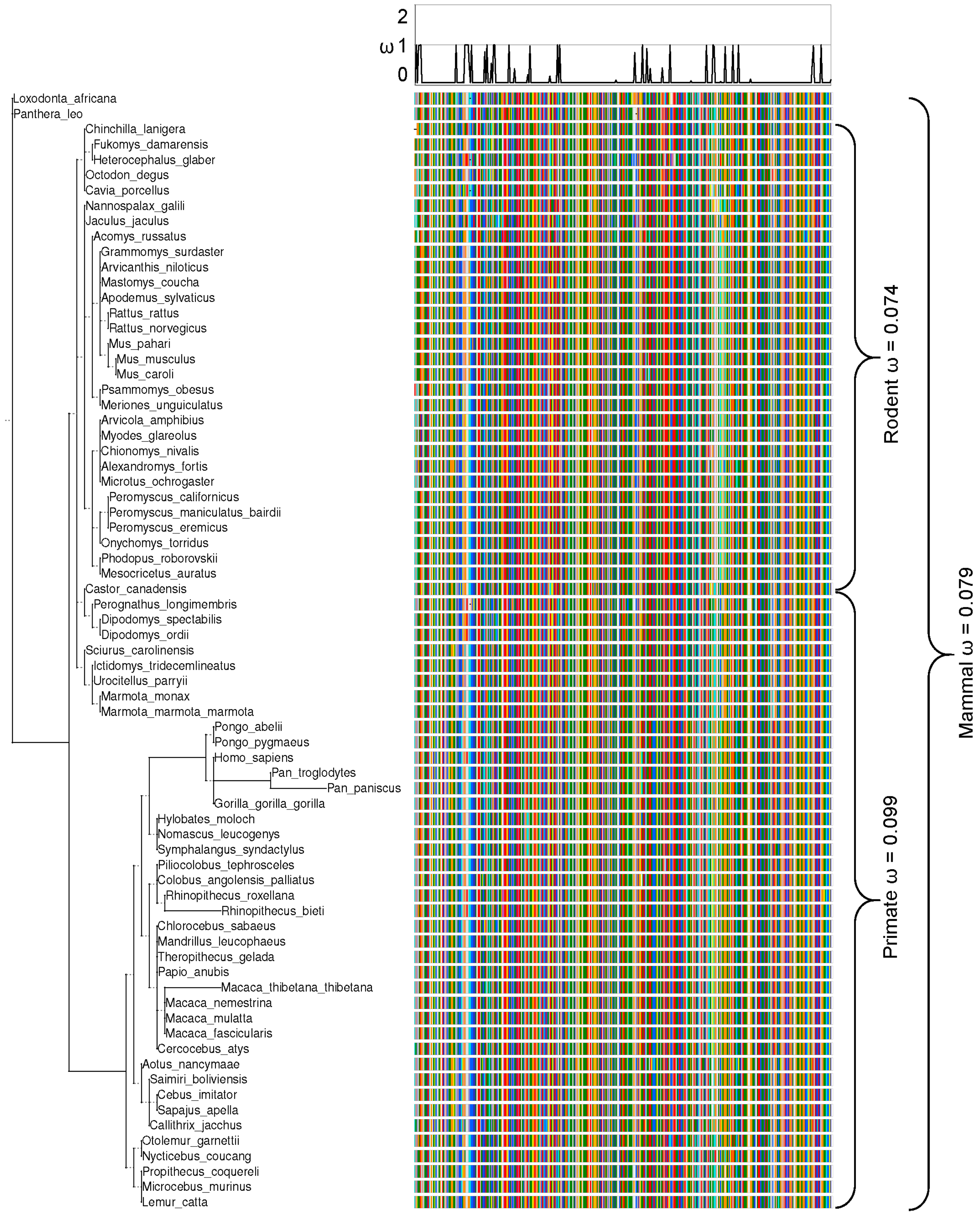


**Figure S1: *Actrt2* is evolutionarily conserved**
Phylogeny of included species, with branch length denoting the number of nucleotide substitutions per codon, and a schematic depiction of the ACTRT2 amino acid aligment utilized in the PAML CodeML analysis. Evolutionary rate (ω) obtained by CodeML model M0 is shown for the whole alignment (Mammal ω) and the primate (Primate ω) and rodent clade (Rodent ω). The graph at the top depicts ω per codon site across the whole tree (CodeML model M2a). See also: Table S1.

#### Figure S2

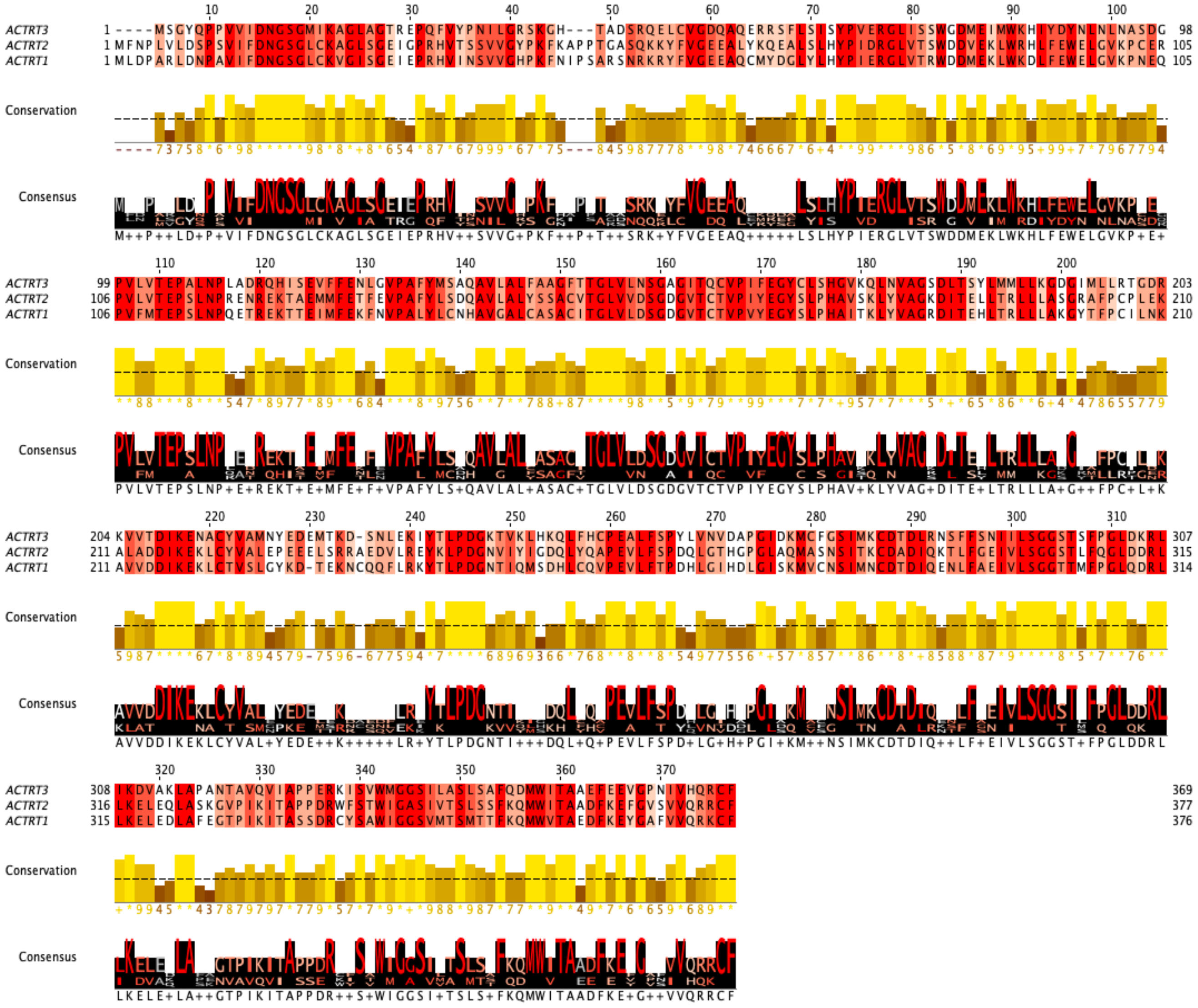

**Figure S2: *Actrt2* shares sequence similarities with other ACTRTs**
Multiple sequence alignment of murine ACTRT1, ACTRT2, and ACTRT3 generated using Jalview. Identical residues are highlighted in red, and similar residues in shades of orange according to conservation. Significantly different residues are shown in white. The conservation histogram (yellow bars) depicts residue-wise conservation scores across the alignment where the bar hight corresponds to the conservation score with higher bars representing increased conservation. The consensus sequence is shown in the lowest bar, with larger letters indicating higher conservation and symbols denoting physicochemical similarity.

#### Figure S3

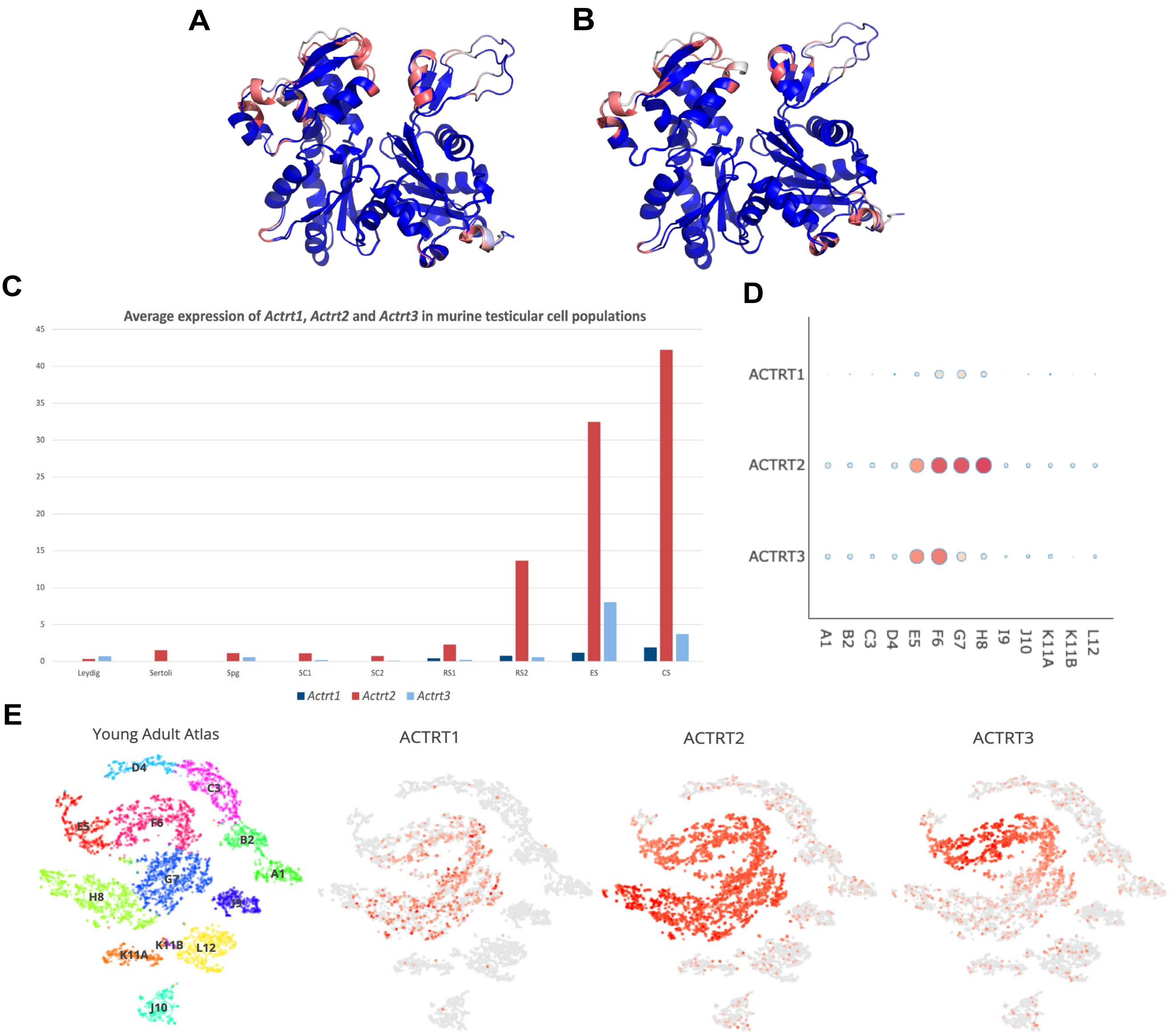


**Figure S3: *Actrt2* shares sequence similarities and expression patterns with other ACTRTs**

A) Structural alignment and confidence mapping of ACTRT1 and ACTRT2 generated using AlphaFold and visualized in PyMOL. Per-residue confidence (pLDDT) is depicted with a colour gradient with blue indicating very high confidence (pLDDT > 90), cyan showing high confidence (70–90) and pink showing low confidence (< 50). The aligned structures exhibit overall high confidence across the core regions, with lower confidence predominantly localized to flexible or exposed regions.

B) Structural alignment and confidence mapping of ACTRT1 and ACTRT3 generated using AlphaFold and visualized in PyMOL. Per-residue confidence (pLDDT) is depicted with a colour gradient with blue indicating very high confidence (pLDDT > 90), cyan showing high confidence (70–90) and pink showing low confidence (< 50). The aligned structures exhibit overall high confidence across the core regions, with lower confidence predominantly localized to flexible or exposed regions.

C) Average expression of *Actrt1*, *Actrt2* and *Actrt3* in cell populations within adult mouse testis: Spg – spermatogonia; SC1 – spermatocytes type 1; SC2 – spermatocytes type 2; RS1 – early round spermatids; RS2 – late round spermatids; ES – elongating spermatids; CS – compacted sperm. Data extracted from Lukassen et al., 2018.

D) Average expression of *ACTRT1*, *ACTRT2* and *ACTRT3* in testis cell populations extracted from Human Testis Atlas. Red indicates high expression while gray indicates low or no expression. Similarly, dot color and size in the dot plot are related to the average expression. A1 – spermatogonial stem cells; B2 – differentiating spermatogonia; C3 – early spermatocytes; D4 – late spermatocytes; E5 – round spermatids; F6 - elongated spermatids; G7 – early sperm cells; H8 – sperm cells; I9 – macrophages; J10 – Endothelia; K11A – Myoid; L12 – Leydig cells. Data extracted from Guo et al. 2018.

E) Average expression of *ACTRT1*, *ACTRT2* and *ACTRT3* in testis cell populations extracted from Human Testis Atlas. Clusters of cells are shown in the left panel: A1 – spermatogonial stem cells; B2 – differentiating spermatogonia; C3 – early spermatocytes; D4 – late spermatocytes; E5 – round spermatids; F6 - elongated spermatids; G7 – early sperm cells; H8 – sperm cells; I9 – macrophages; J10 – Endothelia; K11A – Myoid; L12 – Leydig cells. Data extracted from Guo et al. 2018.

#### Figure S4

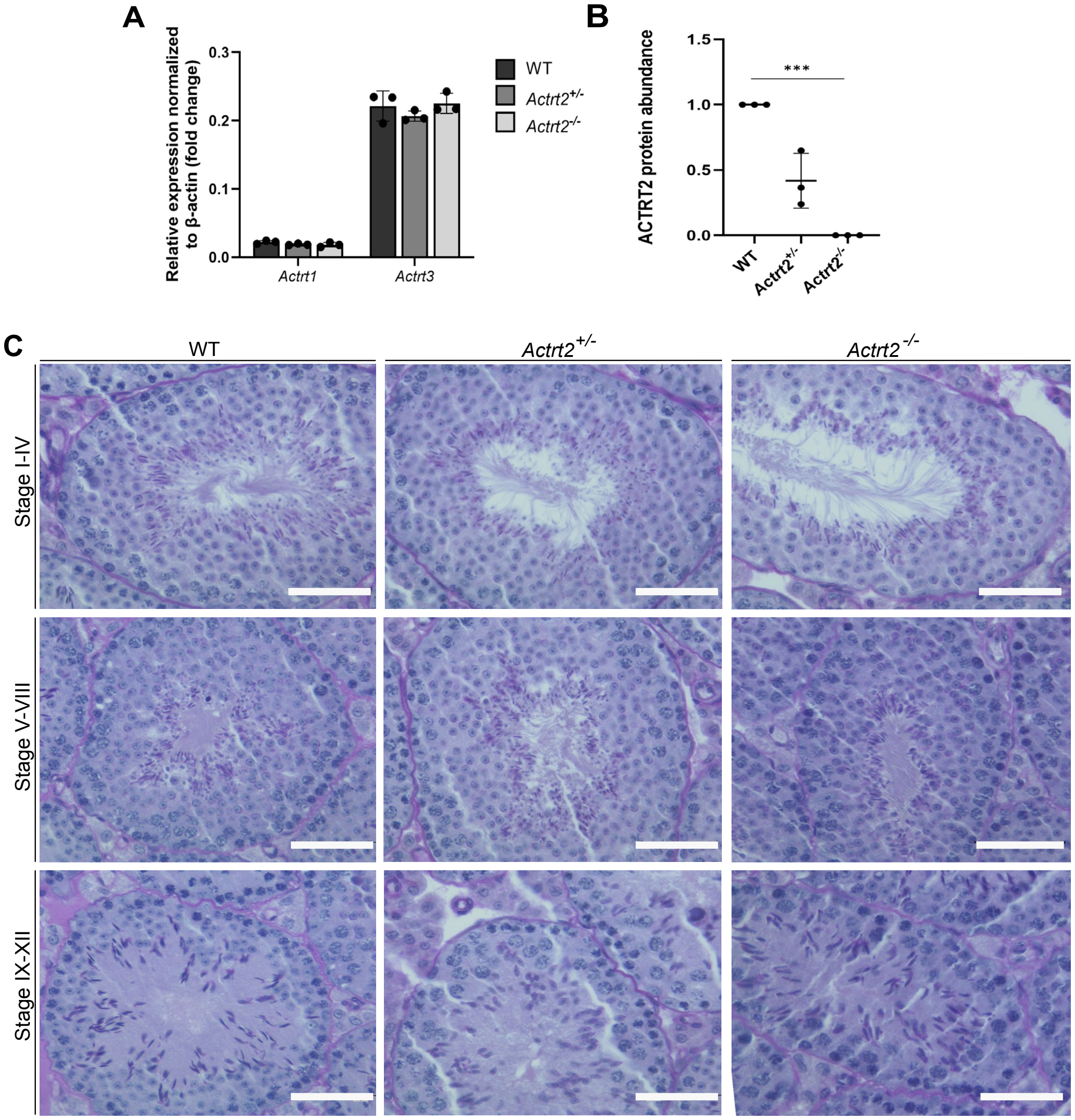


**Figure S4: *Actrt2*-deletion does not affect testicular tissue architecture**

A) Relative expression of *Actrt1* and *Actrt3* in testis tissue from WT, *Actrt2+/-* and *Actrt2-/-* mice analyzed by qRT-PCR and normalized to beta-actin. Analysis was performed using a biological replicate of 3.

B) Quantification of ACTRT2 protein levels in testicular protein lysates from WT, *Actrt2*+/- and *Actrt2-/-* mice analysed by Western blot and normalized to alpha-Tubulin. n=3. ***P<0.001 (one-way ANOVA with Bonferroni correction)

C) PAS staining of testicular tissue sections from WT, *Actrt2*+/- and *Actrt2*-/- mice. Scale bar: 50 µm. Staining was performed on three animals per genotype.

### Figure S5

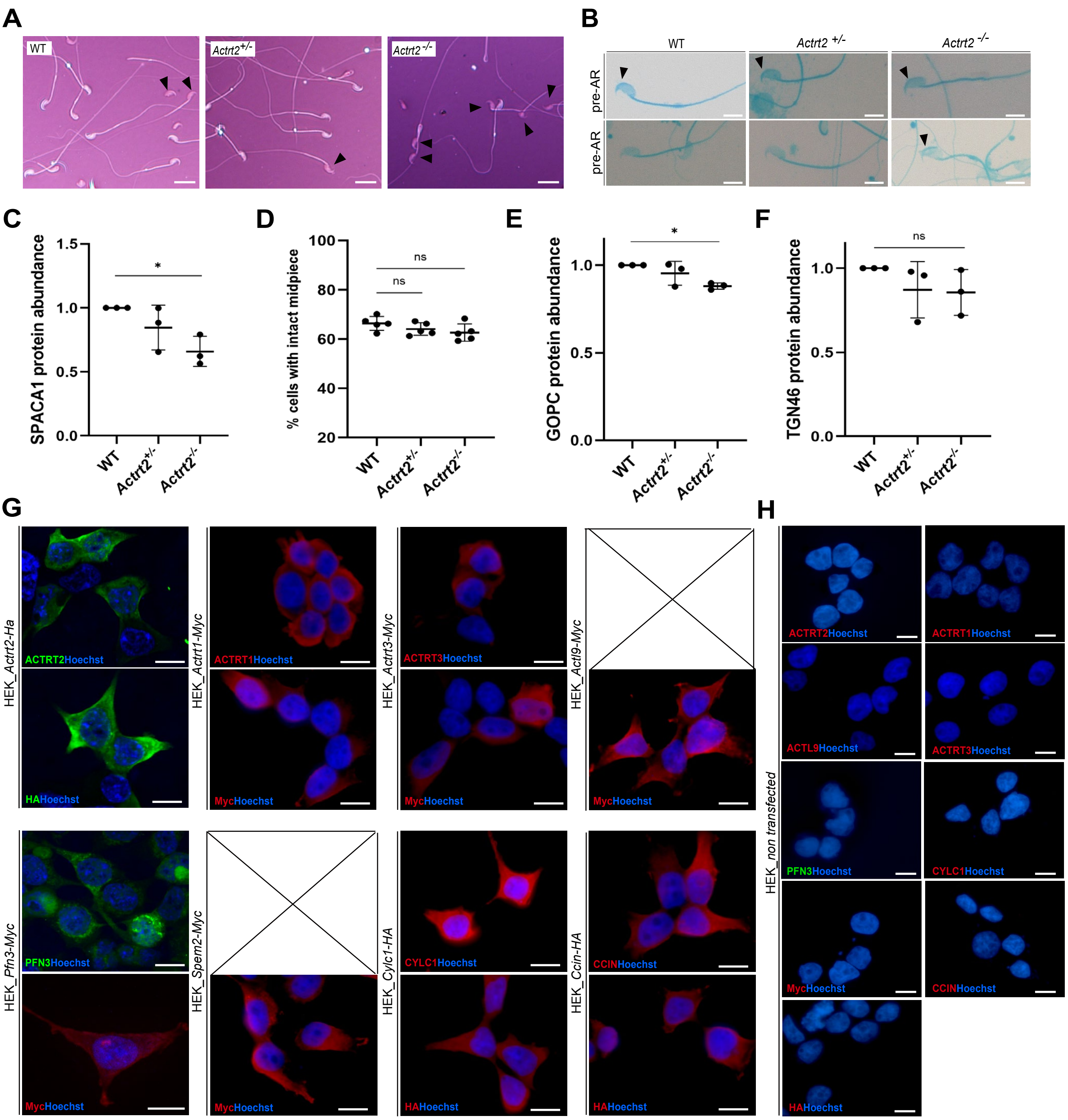


**Figure S5:**

A) Representative images of Eosin-Nigrosine staining of epididymal sperm from WT, *Actrt2+/-* and *Actrt2-/-* male mice. Cells stained pink counted as EN-positive are depicted with black arrowheads. Scale bar: 20 µm.

B) Representative photographs of WT, Actrt2+/- and Actrt2-/- epididymal sperm stained with Commassie blue pre-and post-acrosome reaction. Black arrowheads depict intact acrosomes. Scale bar: 10 µm

C) Quantification of SPACA1 protein levels in testicular protein lysates from WT, *Actrt2+/-* and *Actrt2-/-* mice analysed by Western blot and normalized to alpha-Tubulin. n=3. *P<0.05 (one-way ANOVA with Bonferroni correction)

D) Quantification of sperm cells with midpiece abnormalities observed by MITOred staining in WT, *Actrt2+/-* and *Actrt2-/-* male mice. Black dots represent mean values obtained for each animal included in the analysis. Columns represent mean values ± s.d. (n=5). ns: not significant (two-tailed, unpaired Student's t-test).

E) Quantification of GOPC protein levels in testicular protein lysates from WT, *Actrt2+/-* and *Actrt2-/-* mice analysed by Western blot and normalized to alpha-Tubulin. n=3. *P<0.05 (one-way ANOVA with Bonferroni correction)

F) Quantification of TGN46 protein levels in testicular protein lysates from WT, *Actrt2+/-* and *Actrt2-/-* mice analysed by Western blot and normalized to alpha-Tubulin. n=3. ns: not significant (one-way ANOVA with Bonferroni correction)

G) Immunofluorescent staining of transfected HEK293T cells against ACTRT2, ACTRT1, ACTRT3, ACTL9, PFN3, SPEM2, CYLC1, CCIN, Myc and HA-tag. Nuclei were counterstained with Hoechst (blue). Scale bar: 10 µm.

H) Immunofluorescent staining of non-transfected HEK293T cells against ACTRT2, ACTRT1, ACTRT3, ACTL9, PFN3, SPEM2, CYLC1, CCIN, Myc and HA-tag. Nuclei were counterstained with Hoechst (blue). Scale bar: 10 µm.

#### Figure S6

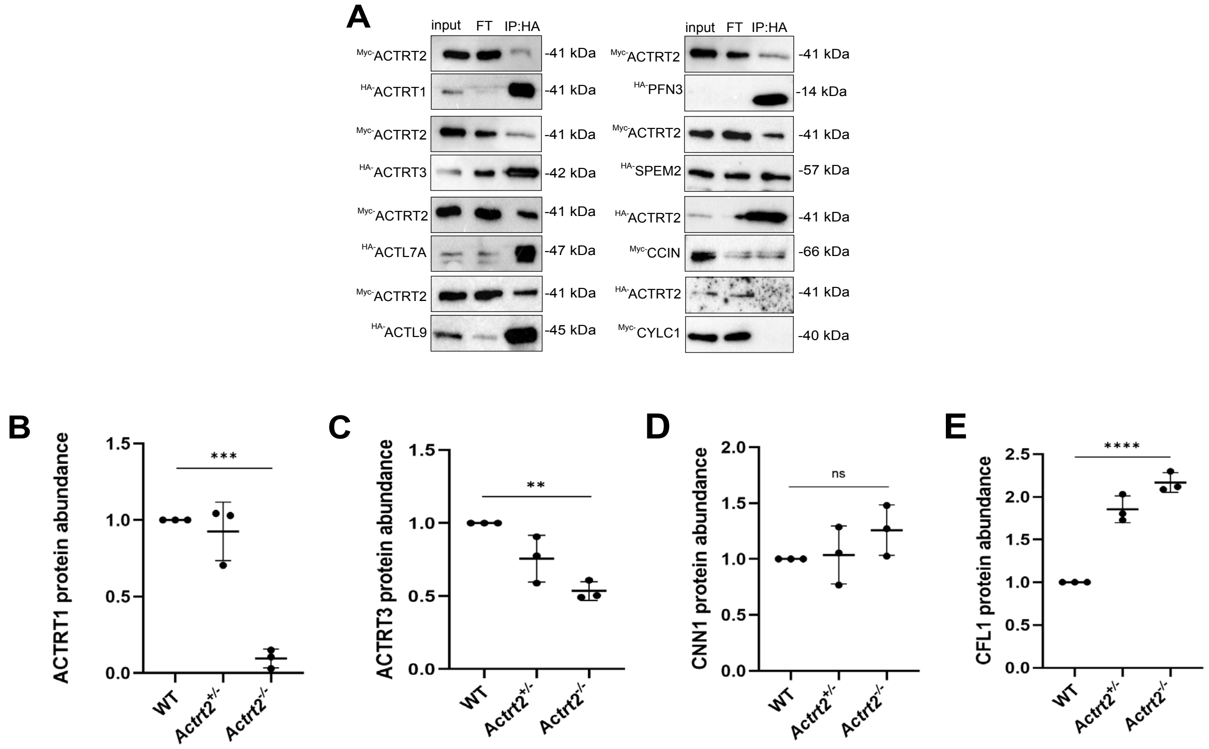


**Figure S6: *ACTRT2* is a structural component of the PT**

A) Co-IP assay of depicting interaction between Myc-ACTRT2 with HA-ACTRT1, HA-ACTRT3, HA-ACTL7A, HA-ACTL9, HA-PFN3 and HA-SPEM2 and the interaction between HA-ACTRT2 with Myc-CCIN. Interaction between HA-ACTRT2 and Myc-CYLC1 was not detected.

B) Quantification of ACTRT1 protein levels in testicular protein lysates from WT, *Actrt2*+/- and *Actrt2-/-* mice analysed by Western blot and normalized to alpha-Tubulin. n=3. ***P<0.001 (one-way ANOVA with Bonferroni correction)

C) Quantification of ACTRT3 protein levels in testicular protein lysates from WT, *Actrt2*+/- and *Actrt2-/-* mice analysed by Western blot and normalized to alpha-Tubulin. n=3. **P<0.01 (one-way ANOVA with Bonferroni correction)

D) Quantification of CNN1 protein levels in testicular protein lysates from WT, *Actrt2*+/- and *Actrt2-/-* mice analysed by Western blot and normalized to alpha-Tubulin. n=3. ns: not significant (one-way ANOVA with Bonferroni correction)

E) Quantification of CFL1 protein levels in testicular protein lysates from WT, *Actrt2*+/- and *Actrt2-/-* mice analysed by Western blot and normalized to alpha-Tubulin. n=3. ****P<0.0001 (one-way ANOVA with Bonferroni correction)

#### Figure S7

A

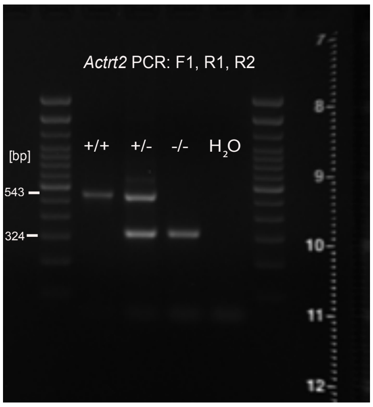


Uncropped picture of the agarose gel depicting genotyping PCR of *Actrt2*-deficient mouse line shown in **Figure 1 E**.

B

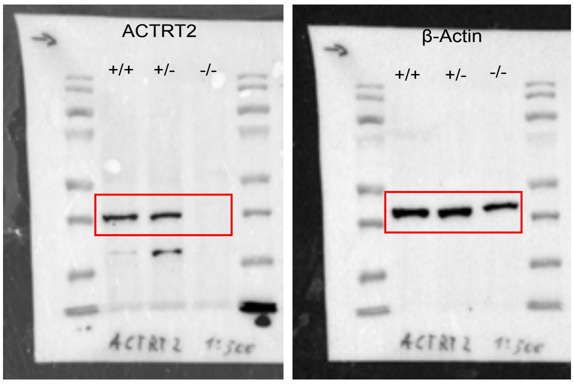


Uncropped picture of the WB membrane incubated with ACTRT2 (left) and β-Actin (right) shown in **Figure 1 H**.

C

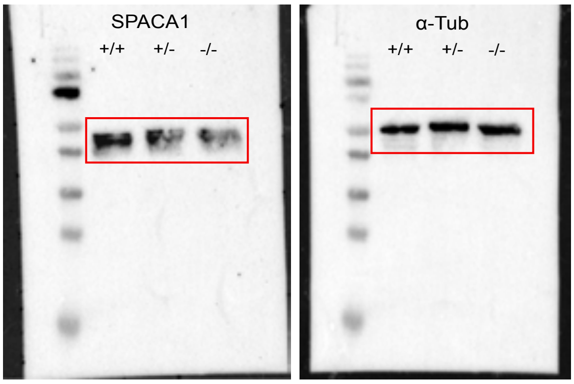


Uncropped picture of the WB membrane incubated with SPACA1 (left) and α-Tubulin (right) shown in **Figure 3 C**.

D

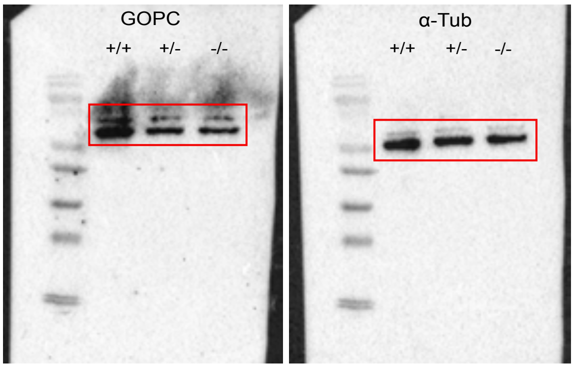


Uncropped picture of the WB membrane incubated with GOPC (left) and α-Tubulin (right) shown in **Figure 4 C**. Membrane was re-blocked before incubating with α-Tubulin.

E

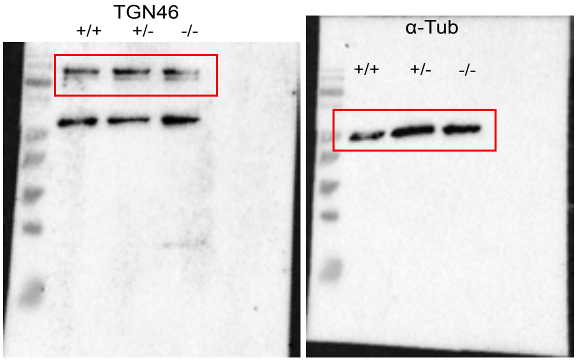


Uncropped picture of the WB membrane incubated with TGN46 (left) and α-Tubulin (right) shown in **Figure 4 C**.

**F**

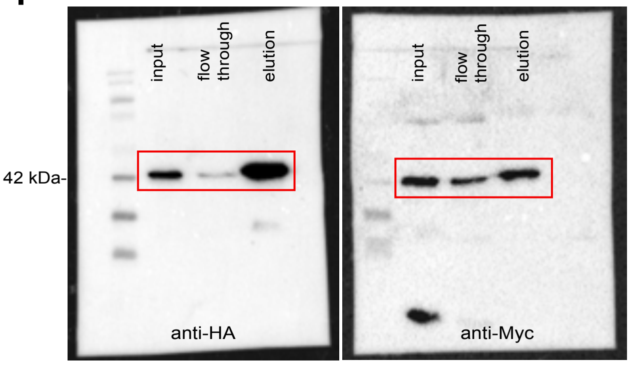


Uncropped picture of the WB membrane depicting Co-IP on HEK cells expressing *Actrt2*-HA and *Actrt1*-Myc using Co-IP kit against HA. Shown in **Figure 5A**.

**G**

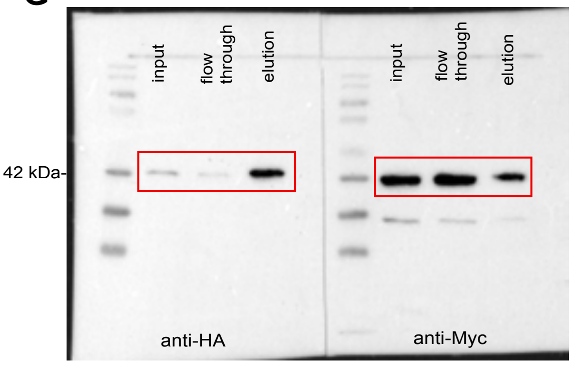


Uncropped picture of the WB membrane depicting Co-IP on HEK cells expressing *Actrt2*-HA and *Actrt3*-Myc using Co-IP kit against HA. Shown in **Figure 5A**.

**H**

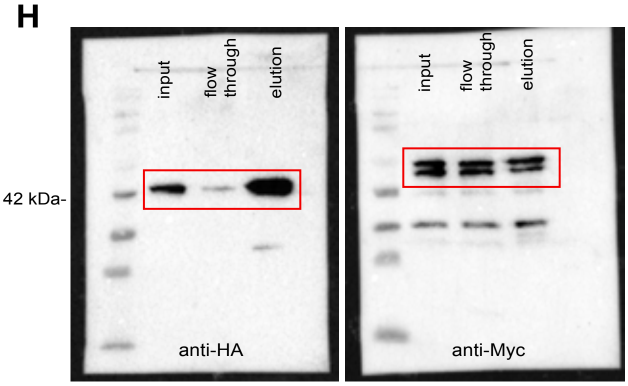


Uncropped picture of the WB membrane depicting Co-IP on HEK cells expressing *Actrt2*-HA and *Actl7a*-Myc using Co-IP kit against HA. Shown in **Figure 5A**.

**I**

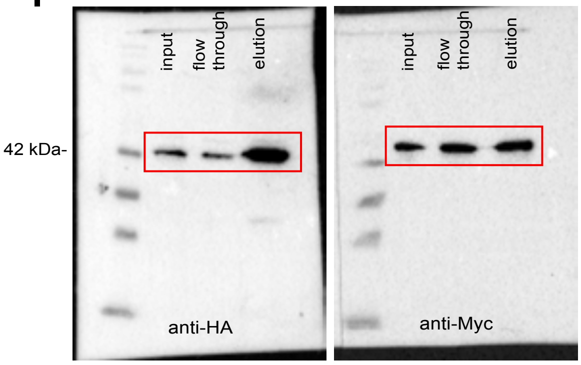


Uncropped picture of the WB membrane depicting Co-IP on HEK cells expressing *Actrt2*-HA and *Actl9*-Myc using Co-IP kit against HA. Shown in **Figure 5A**.

**J**

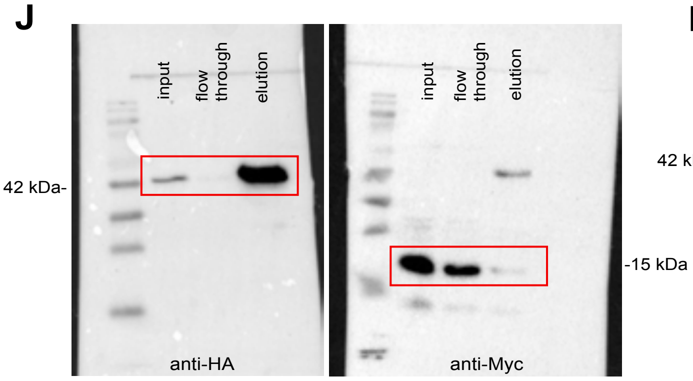


Uncropped picture of the WB membrane depicting Co-IP on HEK cells expressing *Actrt2*-HA and *Pfn3*-Myc using Co-IP kit against HA. Shown in **Figure 5A**.

**K**

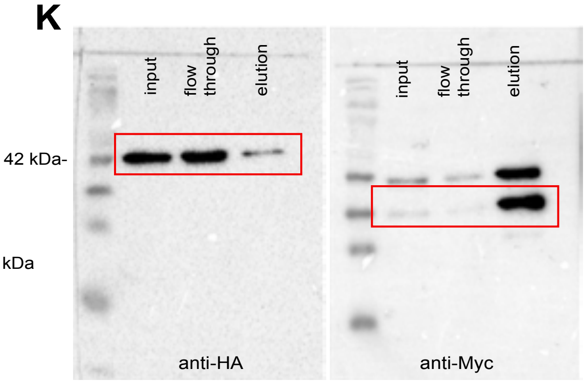


Uncropped picture of the WB membrane depicting Co-IP on HEK cells expressing *Actrt2*-HA and *Spem2*-Myc using Co-IP kit against HA. Shown in **Figure 5A**.

**L**

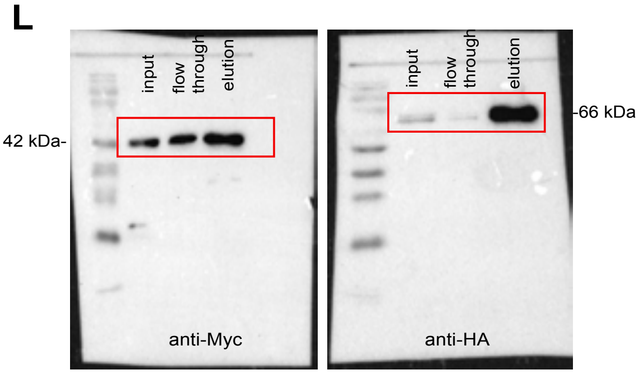


Uncropped picture of the WB membrane depicting Co-IP on HEK cells expressing *Actrt2*-Myc and *Ccin*-HA using Co-IP kit against HA. Shown in **Figure 5A**.

**M**

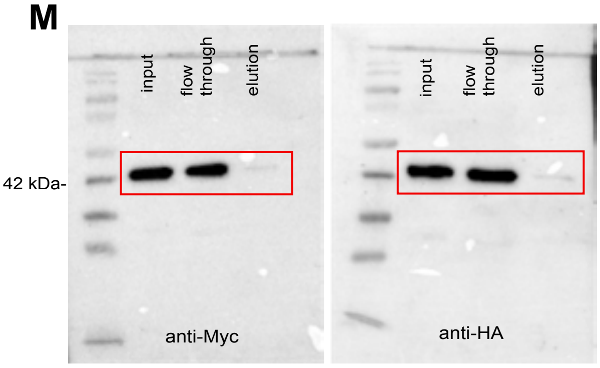


Uncropped picture of the WB membrane depicting Co-IP on HEK cells expressing *Actrt2*-Myc and *Cylc1*-HA using Co-IP kit against HA. Shown in **Figure 5A**.

**N**

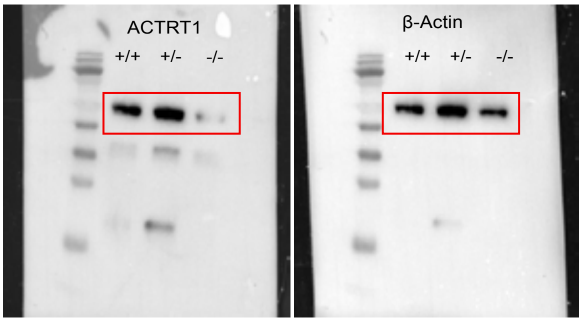


Uncropped picture of the WB membrane incubated with ACTRT1 (left) and β-Actin (right) shown in **Figure 5 C**.

**O**

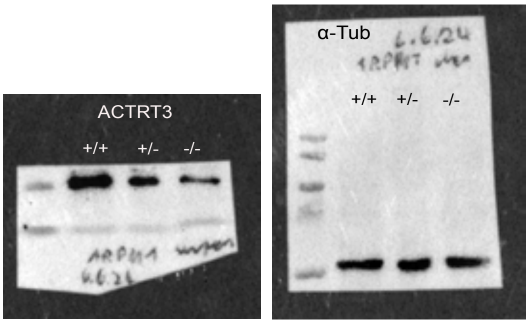


Uncropped picture of the WB membrane incubated with ACTRT3 (left) and α-Tubulin (right) shown in **Figure 5 D**. Membrane was cut to allow simultaneous incubation with two different antibodies in different blocking solutions.

**P**

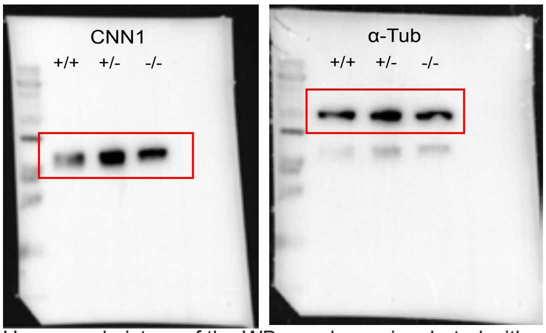


Uncropped picture of the WB membrane incubated with CNN1 (left) and α-Tubulin (right) shown in **Figure 5 C**.

**Q**

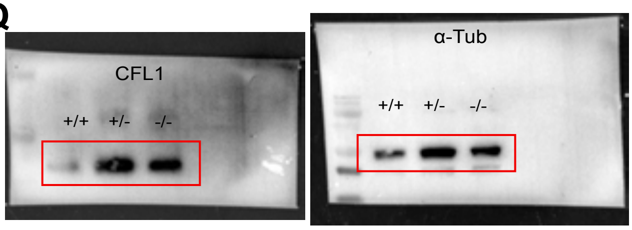


Uncropped picture of the WB membrane incubated with CFL1 (left) and α-Tubulin (right) shown in **Figure 5 D**. Membrane was cut to allow simultaneous incubation with two different antibodies in different blocking solutions.

**Table S1: Protospacer sequences used for generation of *Actrt2*-deficient mouse line**

| Name | protospacer sequence (5´-3´) |
|---|---|
| *Mm.Cas9.Actrt2.sg1* | GGACCAGTTCGTGTCACAAT |
| *Mm.Cas9.Actrt2.sg2* | CAAAGAGTTTGGAGTGTCCG |

**Table S2: Genotyping PCR primer sequences**

| | 5´-3´ | expected band size |
|---|---|---|
| *Actrt2_fwd* | GTCTGCCACGGTCAAGTGTA | WT: 543 bp |
| *Actrt2_rev* | CATACTGGGCAACCACCCTT | KO: 324 bp |
| *Actrt2_int* | AGAGTACAAACTGCCCGACG | |

**Table S3: Antibodies used in this study**

| Antibody | Manufacturer | Catalogue number | Dilution IF | Dilution WB |
|---|---|---|---|---|
| ACTL7A | Proteintech | 17355-I-AP | 1:500 | 1:500 |
| ACTRT1 | Sigma Aldrich | HPA003119 | 1:200 | 1:1000 |
| ACTRT2 | Proteintech | 16992-1-AP | 1:100 | 1:1000 |
| ACTRT3 | Proteintech | 27580-1-AP | 1:500 | 1:1000 |
| CCIN | Progen | GP-SH3 | 1:500 | - |
| CNN1 | Invitrogen | MA5-38095 | - | 1:500 |
| CFL1 | GeneTex | GTX102156 | - | 1:500 |
| CYLC1 | Davids Biotechnology | Custom made (Schneider et al., 2023) | 1:500 | - |
| GOPC | Proteintech | 12163-I-AP | - | 1:500 |
| HA-tag | Proteintech | 81290-1-RR | 1:500 | 1:5000 |
| Myc-tag | Proteintech | 16286-1-AP | 1:500 | 1:1000 |
| PFN3 | Provided by Prof. Dr. Walter Witke | - | 1:250 | - |
| PLCζ1 | Invitrogen | PA5-98556 | 1:250 | - |
| SPACA1 | Abcam | 1007070-2 | - | 1:1000 |
| TGN46 | Invitrogen | MA5-32532 | - | - |
| a-tubulin | Abcam | AB7291 | - | 1:10000 |

**Table S4: qRT primer sequences**

| **Primer name** | **sequence 5´-3´** | **PrimerBank ID (Wang et al. 2012)** |
|---|---|---|
| *Actrt1_fwd* | GAGATTGAACCTCGTCATGTCAT | 13386318a1 |
| *Actrt1_rev* | GCTTCTTCTCCCACAAAGTACCT | 13386318a1 |
| *Actrt2_fwd* | TTTTTGACAACGGATCTGGACTC | 13386316a1 |
| *Actrt2_rev* | TCTTCTGACTAGCTCCCGTGG | 13386316a1 |
| *Actrt3_fwd* | ATGCTCCGGGCATCGACAAG | - |
| *Actrt3_rev* | GGAAGGCAGACAAGGAGGCA | - |
| β-*Actin_fwd* | TGTTACCAACTGGGACGACA | - |
| β-*Actin_rev* | GGGTGTTGAAGGTCTCAAA | - |

**Table S5: Overhang PCR primer sequences**

| **Primer name** | **sequence 5´-3´** |
|---|---|
| *OV_fwd_EcoR1_ACTRT1* | AAAAGAATTCATGCTTGATCCAGCTAGATTAGAT |
| *OV_rev_XhoI_ACTRT1* | TTTTCTCGAGAAAAGCATTTTCTTTGAACCACAAATG |
| *OV_fwd_EcoR1_ACTRT2* | AAAAGAATTCATGTTTAACCCACTGGTACTAGATTC |
| *OV_rev_XhoI_ACTRT2* | TTTTCTCGAGAGAAGCACCGCCTCTGGAC |
| *OV_fwd_ApaI_ACTRT3* | AAAAGGGCCCATGAGCGGCTACCAGCCAC |
| *OV_rev_KpnI_ACTRT3* | TTTTGGTACCAAAGCATCTCTGGTGTACAATGTTGG |
| *OV_fwd_EcoRI_ACTL7A* | AAAAGAATTCATGTCTCTGGATGGTGTGTG |
| *OV_rev_KpnI_ACTL7A* | TTTTGGTACCGAAGCACCTTCTGTAGAGGA |
| *OV_fwd_EcoR1_ACTL9* | AAAAGAATTCATGGATGTCAATGGACACCCAAAG |
| *OV_rev_Kpnl_ACTL9* | TTTTGGTACCGTAGCATTTTCGGTATACAACCTG |
| *OV_fwd_BglII_CCIN* | AAAAGGTACCATGAAACTGGAATTCACAGAGAAA |
| *OV_rev_Kpnl_CCIN* | TTTTGGTACCAATTCTTTGAAGAATCTTGCAAGG |
| *OV_fwd_NcrI_CYLC1* | AAAAAGATCTATGTCTCTTTCAAAACTGGACAGTG |
| *OV_rev_KpnI_CYLC1* | TTTTGGTACCGAGCAGTTTATGAATCCACTCCA |
| *OV_fwd_EcoRI_PFN3* | AAAAGAATTCATGAGTGACTGGAAGGGCTACATCAGTGC |
| *OV_rev_KpnI_PFN3* | TTTTGGTACCAGAGCACTGCTCACGCAGCCCACCAA |
| *OV_fwd_EcoRI_SPEM2* | AAAAGAATTCATGGAAAACCAGCTGTGGCAGAACA |
| *OV_rev_XhoI_SPEM2* | TTTTCTCGAGAGTTAAGTTTCCCTGTCCGGCTTCTA |

**Table S6: Results of the evolutionary analysis using CodeML (PAML4.9)**

| Selection on: | Gene / Clade | | LnL null model | LnL alternative model | LRT | p | ω | interpretation | | | | |
|---|---|---|---|---|---|---|---|---|---|---|---|---|
| whole sequence, whole tree | *Actrt2* | | 16217.474 (M0) | 18371.069 (M0fix) | -4307.19 | n.s. | 0.079 (M0) | overall highly conserved | | | | |
| whole sequence, clades / lineages | *Actrt2* | Primates | 16217.474 (M0) | 16211.556 (MC) | 29.09 | >0.01 | 0.099 (MC) | both lineages highly conserved | | | | |
| | | Rodentia | | | | | 0.074 (MC) | | | | | |
| codon sites, whole tree | *Actrt2* | | 15975.770 (M1a) | 15975.770 (M2a) | 0 | n.s. | n.a. | **prop 0** | **prop 1** | **prop 2a** | **PSS** | **PUR** |
| | | | | | | | | 0.91 | 0.08 | n.a. | n.a. | 89% of sequence |

LnL = log Likelihood value of the model; LRT = Likelihood Ratio Test; p = p-value of the LRT; ω = evolutionary rate (dN/dS); prop = proportion of sites assigned to the respective site class; PSS = significantly positively selected sites; PUR = sites under significant purifying selection / conserved sites; for explanation of the models see M&M.
